# Mesenchymal Progenitors set the homeostatic inflammatory milieu via the TAK1-NFkB axis

**DOI:** 10.1101/2022.12.11.519940

**Authors:** Theret Marine, Messing Melina, White Zoe, Tung Lin Wei, Rempel Lucas, Hamer Mark, Hashimoto Joshua, Li Fangfang, Brasssar Julyanne, Li Yicong, Sauge Elodie, Shin Samuel, Day Katie, Uppal Manjosh, Low Marcela, Eisner Christine, Shintaro Sato, Shizuo Akira, Hughes Michael, Bernatchez Pascal, Kelly M McNagny, Fabio M.V. Rossi

## Abstract

The ability of mesenchymal stromal cells to modulate inflammation is at the basis of the ongoing interest in their therapeutic potential. Yet, reliable success in clinical trials is limited, possibly due to a limited understanding of their impact on the inflammatory milieu in physiological conditions. Here we show that, at steady state, mesenchymal progenitors regulate the balance between type 1 and type 2 inflammatory milieus by acting on innate immune cells through the TAK1-NFkB pathway. Suppressing the constitutive activity of this pathway in MPs leads to skewing of the immune system toward systemic Type 2 inflammation (Th2). These changes have significant effects on diseases with an important inflammatory component, leading to a worsening of disease in a preclinical model of Th2-dependent Asthma, and a reduction of symptoms associated with Th1/Th17-dependent experimental autoimmune encephalitis.

## Introduction

Mesenchymal progenitors (MPs) are present in most, if not all, the tissues of the body where they fulfil a variety of essential functions ^1^. While MPs are known to maintain tissue homeostasis and architecture by producing extracellular matrix (ECM) proteins, their capacity to orchestrate inflammation has been highlighted during development, postnatal growth, adulthood ^2,3^. However, recent in vivo data suggests that these cells can polarize inflammation in opposite directions depending on the environment^4^, secreting chemokines that attract typical type 1 cells such as neutrophils and inflammatory monocytes in response to acute damage^5,6^, and typical type 2 cells such as regulatory T cells (Tregs), eosinophils, and ILC2s in chronic disease^7–10^. This dynamic immunomodulatory activity may help rationalize failed cell-based therapies if improperly regulated upon transplantation ^11,12^. Polarization of mesenchymal cells to a type1 pro-inflammatory state has been linked to the activation of TLR family members^13^. Transforming growth factor β-activated kinase 1 (TAK1) is a central kinase downstream of multiple inflammatory cytokines including transforming growth factor β (TGFβ), interleukin 1β (IL1β), tumor necrosis factor α (TNFα), as well as TLRs ^14–17^. Downstream of these receptors, Tak1 is a key intermediate in the signalling pathway leading to the activation of NFkB. Thus, TAK1 has been extensively studied in the context of innate inflammation within immune cells in order to dampen inflammatory reactions ^18^. In contrast, its deletion in other cell types such as mouse embryonic fibroblasts, hepatocytes, keratinocytes, or enterocytes has been shown to induce cell death and inflammation ^19–23^. This led us to hypothesize that TAK1 could have an important role in MP inflammatory functions. Here we explore the role of TAK1 in mesenchymal progenitor cells and show that in its absence, the systemic inflammatory environment is strongly biased towards Th2 responses. We further show that this affects the severity of inflammatory diseases, increasing it or decreasing it depending on the role that inflammation plays in their pathogenesis. Thus, modulating MP immune function impacts the outcome of inflammatory diseases, and may represent a strategy to make the benefit of MP cell therapies more reliable.

## Results

### HIC1^ΔTAK1^ mice display a systemic type 2 inflammatory response

We deleted TAK1 specifically in MPs by exploiting a new mouse strain in which CRE-ERT2 is driven by Hypermethylated in Cancer 1: Hic1 ^7,24^ (Figure 1A). To explore potential changes in circulating cells, splenocytes were harvested and processed for mass cytometry (CyTOF) with a panel of 21 antibodies. 42 phenotypic clusters were identified (Figure 1A, S1A-B, Table S1) and were broadly labelled as B cells, CD4 T cells, CD8 T cells, NK cells, eosinophils, myeloid cells, and type 2 innate lymphoid cells (ILC2) (Figure S1B, Table S1). When comparing TAK1^ΔHIC1^ mice to their littermate controls TAK1^WT^, we noticed a 2-fold (p<0.001) increase in eosinophils as well as a striking 34-fold (p<0.05) increase in ILC2s (Figure S1G and I). We also noted a significant decrease in B cells and NK cells, but no changes in T cells and myeloid cells (Figure S1C-F, H). Elevated levels of circulating eosinophils and ILC2s were then validated by flow cytometry on freshly harvested blood (Respectively: +219%, p<0.0001, Figure 1C and S2A; +348%, p<0.0001, Figure 1D and S2B) and in various tissues (Figure 1E and F, Figure S2C and D). To further evaluate the inflammatory milieu in the blood of HIC1^ΔTAK1^ mice, cytokines present in the serum were quantified by mesoscale assay. IL-15, IL-5, IL-4, GM-CSF, IL-6, IL-16, and EPO were all elevated in the serum of HIC1^ΔTAK1^ mice (Figure S1J, Table S2). Concomitantly, serum IgE was increased by 2.7fold (p<0.01, Figure 1G). These results suggest that HIC1^ΔTAK1^ mice display a predominantly Type 2 systemic inflammatory response with B cell activation. While it has been previously shown that a sub-population of CD4T cells expressed Hic1^25^, the analysis of cytokine expression in splenic T cells, as well as the proportion of Tregs in the mesenteric lymph nodes (mLN) did not reveal evidence of T cell activation nor suggested skewing toward Type 2 inflammation (Figure S3A-D). Consistent with a lack of T cell involvement, blood eosinophilia was still found in HIC1^ΔTAK1^ mouse model lacking T and B cells (RAG1-KO, Figure S4A, +220%, p<0.001). In contrast, we found a dramatic increase in the frequency of KLRG1+ cells within circulating ILC2 (+461%, p<0.001, Figure 1H), as well as increased production of IL-4, IL-13, and IL-5 (Figure 1I, S4B) within these cells, suggesting they adopted an activated state. In summary, the changes we observed in HIC1^ΔTAK1^ mice at steady state are consistent with a constitutively activated Type 2 inflammatory response that is independent of T cells and induced by ILC2 activation.

**Figure 1:**
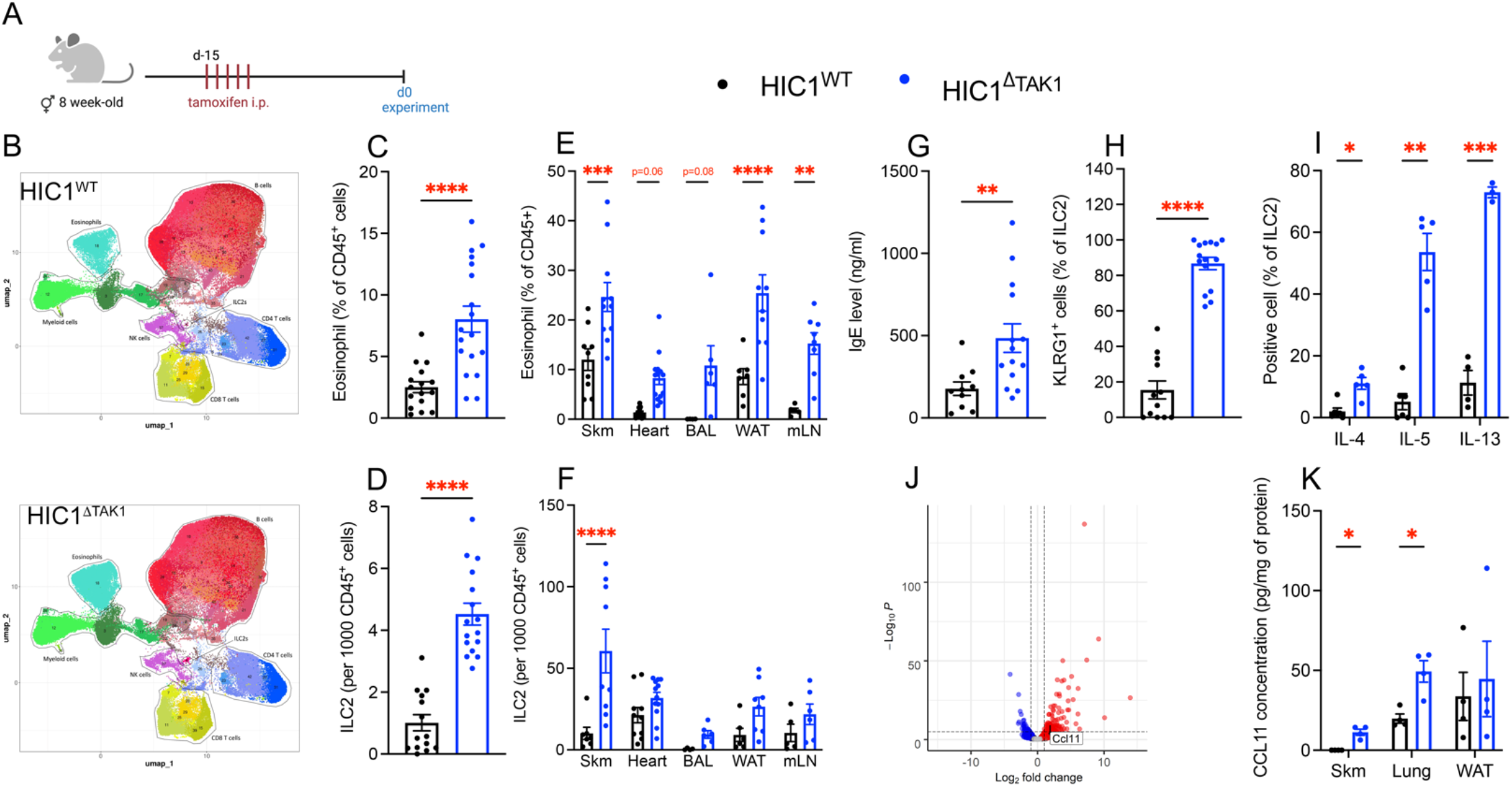
HIC1^ΔTAK1^ mice display a systemically activated type 2 inflammatory response. After tamoxifen induction (A), splenocytes were isolated and run against a 21 antibody CyTOF panel (B; n=3). Eosinophil and ILC2 content were validated on freshly harvested blood (C, D; n=16-18), skeletal muscle (Skm), cardiac muscle (heart), bronchoalveolar lavage (BAL), white adipose tissue (WAT) and mesenteric lymph nodes (mLN) (E, F; n=4-14). Blood IgE levels were measured by ELISA (G; n=10-14) and the percentage of KLRG1+ ILC2 were quantified (H; n=12-15). ILC2 from the spleen were stimulated *in vitro* and the production of IL-4, IL-5, and IL-13 was quantified by flow cytometry (I; n=4-6). RNASeq was performed on Skm-resident MPs and differentially expressed genes are shown as a volcano plot (J; n=3-5). CCL11 concentration was measured on Skm, lung and WAT (K; n=4). HIC1^ΔTAK1^ versus HIC1^WT^: * p<0.05; ** p<0.01; *** p<0.001; **** p<0.0001

### HIC1^ΔTAK1^ mesenchymal progenitors turn on an inflammatory program

To better understand the cellular interaction between MPs, eosinophils and ILC2 in the HIC1^ΔTAK1^ mouse model, skeletal muscle (skm) resident MPs were sorted using a Td.tomato reporter and subjected to RNAseq (Figure S1K). Gene ontology analysis highlighted changes in the inflammatory prolife of HIC1^ΔTAK1^ MPs, with an increase of biological processes associated to inflammation such as “inflammatory response” or “chemotaxis” (Figure S1L). Critically, HIC1^ΔTAK1^ MPs also express higher level of *Ccl11* (eotaxin-1) (Figure 1J Tables S3) which has been shown to play an important role in maintaining healthy eosinophil levels in adipose tissue ^9,26^. While we could not detect changes in circulating CCL11 levels (Figure S1M), its increase at the tissue level was confirmed by ELISA (Figure 1K). Interestingly, CCL11 is produced by white adipose tissue (WAT) tissue resident MSCs in response to IL4/IL-13 stimulation by ILC2 ^9^, cytokines that are up-regulated in our model (Figure S1J and 1I). Altogether, these results suggest that the hyper-eosinophilia is a secondary event due to the increased level of activation of ILC2 and the consequent increase in type2 cytokines in the HIC1^ΔTAK1^ mouse model.

### HIC1^ΔTAK1^ mice exhibit biased hematopoiesis

An increased proportion of circulating eosinophils in the HIC1^ΔTAK1^ mice expressed progenitor marker CD34+, suggesting an activation of the BM progenitor pool (Figure S4C) ^27,28^. MPs are a critical component of the hematopoietic stem cell (HSC) and progenitor niche and are well-known to support hematopoiesis in adulthood ^29^. In the HIC1^ΔTAK1^ mice, we did not see changes in the total number of Lin-Sca1+ cKit+ cells (LSK), a population that contains multiple types of multipotent progenitors (Figure 2A and S5A). However, we found that phenotypic Long Term (LT)-HSC number were decreased by 25% (p<0.05, Figure 2B and S5A) and that BM hematopoiesis was shifted toward granulopoiesis (Figure 2C and S5B), which resulted in increased numbers of eosinophil progenitors (EOP) (+182%, p<0.05, Figure 2D, and S5C). Altogether, we demonstrate that MPs regulate the inflammatory response at a systemic level, mounting a coordinated type 2 response that includes increased production of eosinophils in the bone marrow as well as their recruitment to peripheral tissues. Our data further suggest that a constitutive level of TAK1 activity in MPs is required to maintain balanced hematopoiesis.

**Figure 2:**
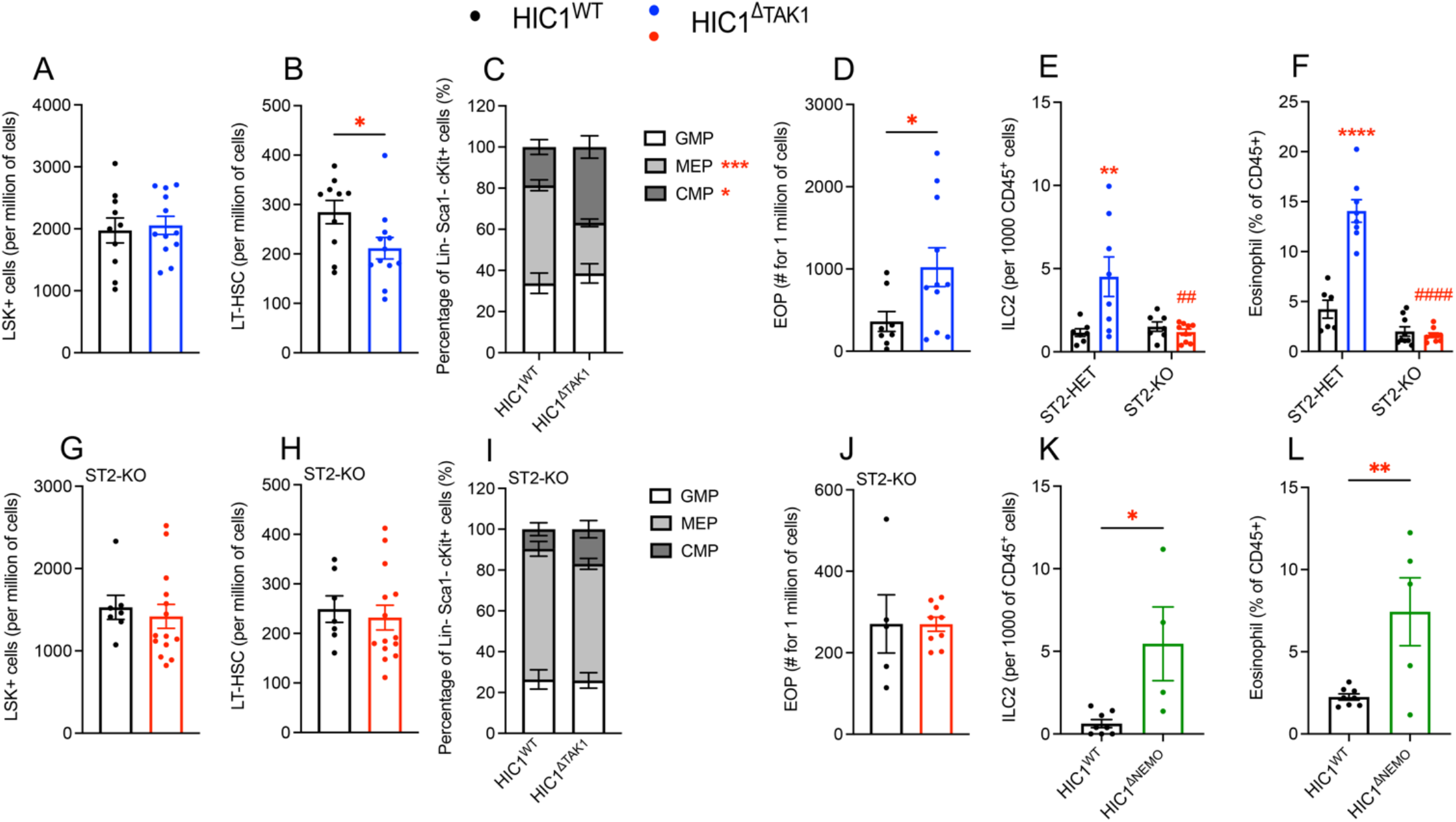
HIC1^ΔTAK1^ phenotype originates from bone marrow, is dependant on ILC2 and involves the NF-κB signaling pathway. Bone marrow from HIC1^WT^ and HIC1^ΔTAK1^ mice was harvested and stained for Lin-Sca+ cKit+ (LSK) cells (A; n=10-12), Long Term-HSC (B; n=10-12) common myeloid progenitor (CMP), granulocyte/monocyte progenitor (GMP), megakaryocyte/erythroid progenitor (MEP) (C; n=6-9), and eosinophil progenitors (EOP) (D) n=8-11. HIC1^ΔTAK1^ mice were bred with ST2-KO mice, and ILC2 (E) and eosinophil (F) content was quantified on freshly harvested blood (n=6-9). Bone marrow was harvested and stained for LSK cells (G; n=7-14), LT-HSC (H; n=7-14), CMP, GMP, MEP (I; n=3-6), and EOPs (J; n=5-9). ILC2 (K) and eosinophil (L) content were quantified on freshly harvested blood of HIC1^ΔTNEMO^ and their controls (n=5-8). HIC1^ΔTAK1^ versus HIC1^WT^: * p<0.05; ** p<0.01; *** p<0.001; **** p<0.0001 ST2^+/-^HIC1^ΔTAK1^ versus ST2^-/-^HIC1^ΔTAK1^ ^T^: ### p<0.001 HIC1^ΔNEMO^ versus HIC1^WT^: * p<0.05; ** p<0.01

### ILC2s are the primary effectors of the TAK1-KO phenotype

ILC2 activation and expansion is often due to an increase in IL-33 or IL-25 ^30,31^. Surprisingly, however, no changes in IL-33 or IL-25 concentrations were noticeable in the blood of HIC1^ΔTAK1^ mice (Figure S1J and S4D, Table S2). Expression of *Il33* was not modified in HIC1^ΔTAK1^ MPs (Table S3) and concentration of IL-33 was also not found increased in various tissues (Figure S4D-E). Moreover, the absence of changes in Tregs suggests that IL-33 is not dysregulated in this system (Figure S3D)^32^. We sought to understand if the blood eosinophilia was due to a primary event of the deletion of TAK1 in MPs, or if it was a consequence of the activation of ILC2s. To do so, we crossed the HIC1^ΔTAK1^ mice with the ST2^-/-^ mouse model, which lack the receptor for IL-33 (IL1RL1) and show impaired ILC2 activation (Figure 2E)^33^. By doing so we prevented blood eosinophilia (Figure 2F), and restored hematopoiesis similar to that observed in ST2^-/-^ :HIC1^WT^ control mice (Figure 2G-J). This suggests that in HIC1^ΔTAK1^ mice, ILC2 activation is required to induce blood hyper-eosinophilia and promote a type2 environment. As previously suggested^34,35^, this takes place in a T cell independent manner.

### Blocking Nuclear factor-κB activation phenocopies the HIC1^ΔTAK1^ phenotype

TAK1 is a pleiotropic kinase with many downstream targets such as extracellular signal-regulated kinase (ERK), p38, c-Jun N-terminal kinases (JNK), or Nuclear factor-κB (NF-κB) (reviewed in ^36^). NF-κB in particular plays a key role in regulating inflammation within immune cells^37^, and a reduction in NF-κB activation in HIC1^ΔTAK1^ mice seems likely to mediate the observed phenotype. To test this, we generated a mouse strain in which the NF-κB pathway is dampened in HIC1+ cells by deletion of the upstream regulator NEMO ^38^ (HIC1^ΔNEMO^) and analyzed blood eosinophil and ILC2 frequencies. We observed the same phenotype as seen in the HIC1^ΔTAK1^ mouse (Figure 2K and L) confirming that the TAK1-NF-κB axis is constitutively required in MPs to maintain a balanced inflammatory environment at steady state.

### Deletion of TAK1 in MPs protects mice against Th1 disease

Experimental autoimmune encephalomyelitis (EAE) is designed to mimic multiple sclerosis (MS) development in mice (Figure 3A). MS is Th1/Th17 driven ^39,40^, and we sought to test whether the Th2 bias of HIC1^ΔTAK1^ mice would result in reduced susceptibility toward this neurodegenerative disease. Indeed, it has been shown previously that promotion of Th2 inflammation delays the appearance of MS symptoms ^41^. At resting state, we found that ILC2s were present in higher numbers throughout the CNS of HIC1^ΔTAK1^ mice (Figure S6A-B). On the contrary only the spinal cord (SP) of HIC1^ΔTAK1^ mice contained more eosinophils compared to control mice (Figure S6C-D). The number of microglia did not change in the brain or the spinal cord (SP) of HIC1^ΔTAK1^ mice compared to controls (Figure S6C, E). After EAE induction, HIC1^ΔTAK1^ mice exhibited delayed disease onset and milder symptoms (Figure 3B-C, S7A). When harvested at the clinical score of 3, brain and SP of HIC1^ΔTAK1^ mice contained similar numbers of ILC2s but more eosinophils than controls (Figure 3D-E, S7B-C). Overall, these results suggest that deletion of TAK1 in MPs can provide protection from Th1/Th17 driven diseases such as EAE.

**Figure 3:**
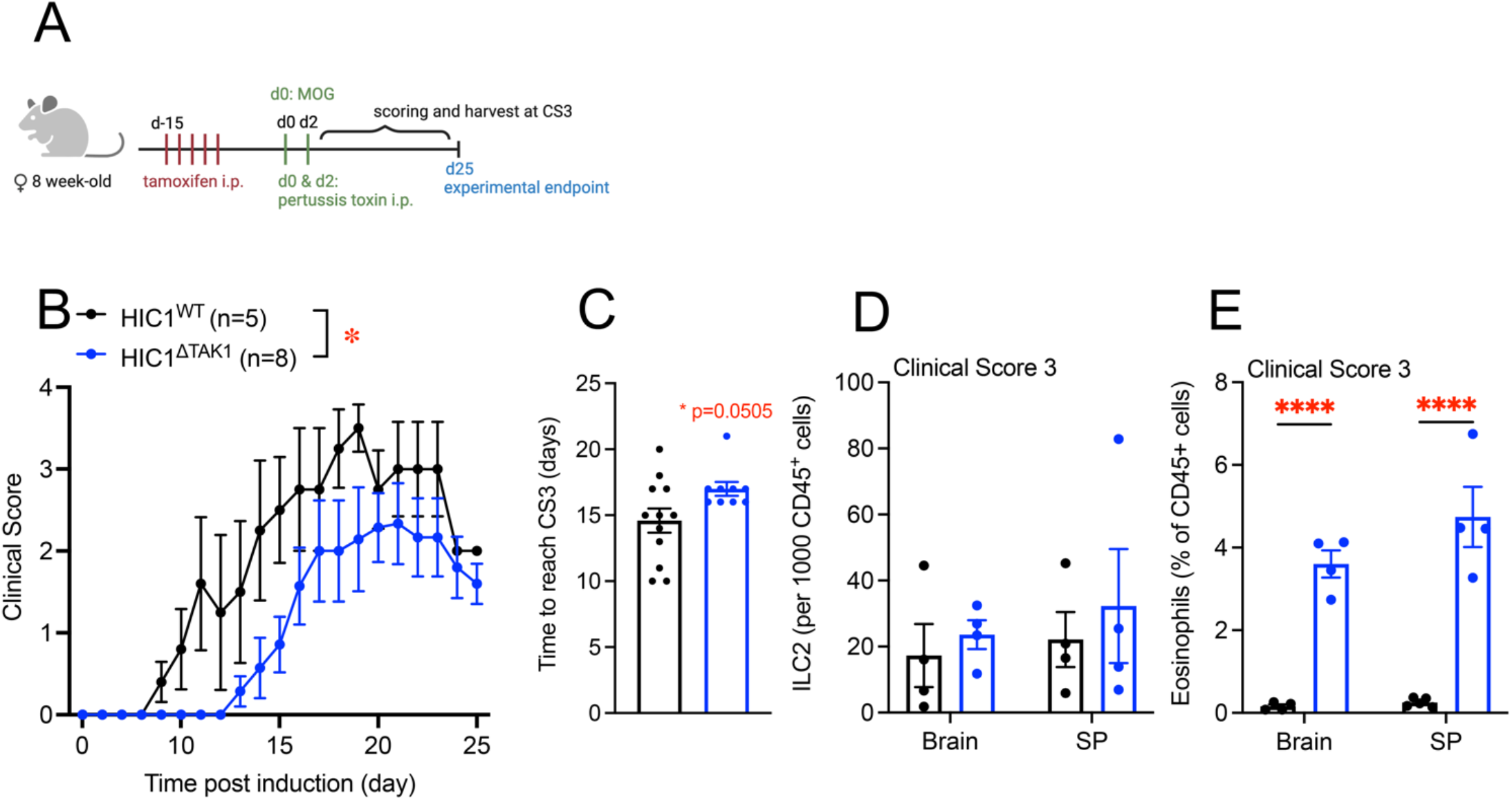
HIC1^ΔTAK1^ mice are protected against experimental autoimmune encephalomyelitis (EAE). After tamoxifen injections, EAE was induced in HIC1^WT^ and HIC1^ΔTAK1^ mice (A). Clinical scores were recorded up to 25 days post-induction (B; n=5-8) and the time to reach clinical score (CS) 3 was calculated (C; n=9-12). Total amounts of ILC2 (D) and eosinophils (E) in both brain and spinal cord (SP) were quantified by flow cytometry in CS3 scored mice (n=4). HIC1^ΔTAK1^ versus HIC1^WT^: * p<0.05; **** p<0.0001

### HIC1^ΔTAK1^ mice are more sensitive to Th2 disease

Next, we investigated whether the systemic Th2 inflammation displayed by HIC1^ΔTAK1^ mice may have harmful consequences for the development of allergic disease, a typical Th2-driven pathology. First, control and HIC1^ΔTAK1^ mice were exposed to papain through inhalation in order to induce a Th2 allergic response (Figure 4A) ^42^ and lung function was analysed. HIC1^ΔTAK1^ lung capacity was drastically decreased after papain damage compared to HIC1^ΔTAK1^ mice exposed to PBS only, or compared to control mice treated with papain (Figure 4B). Reduced lung function was caused by increased lung stiffness (Figure S8A-F) and an increase in inflammation as shown by H&E staining (Figure 4C) resulting in a histological score increase of 44% compared to papain-treated control mice (p<0.05, Figure 4D ^43^. Collagen and mucus deposition were unchanged (Figure S8G and H) suggesting that inflammation was the leading cause of the decrease in lung functions. Indeed, papain-treated HIC1^ΔTAK1^ mice also displayed higher levels of IgE in the blood (Figure S8I), and a higher number of eosinophils in the lung (Figure S9A-E), which confirms over-activation of the Th2 inflammatory response. Strikingly, the loss of ST2 completely rescued the increased loss of lung function in mice lacking TAK1 in MPs (Figure 4E-G, and S10A-H), again demonstrating the importance of this pathway in mediating the effects of MPs toward inflammation

**Figure 4:**
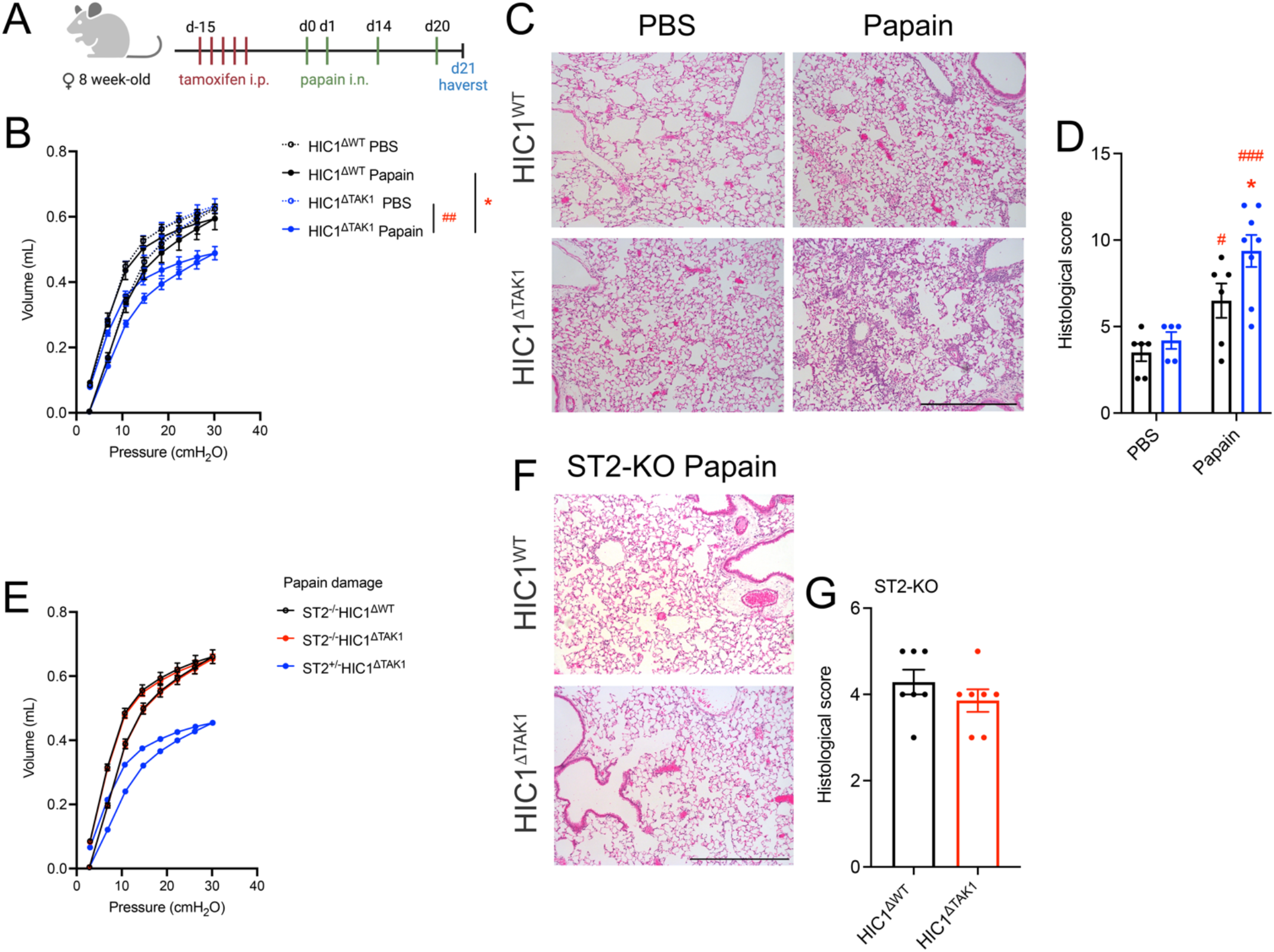
HIC1^ΔTAK1^ mice are a higher sensitivity to the allergen papain. After tamoxifen injections, HIC1^WT^ and HIC1^ΔTAK1^ mice were subjected to intra-nasal papain injections to induce an allergic reaction (A). Lung function was measured using Flexivent® (B; n=5-7) and lung morphology was histologically assessed (C) and scored (D) by Hematoxylin and Eosin Staining (H&E; n=7). HIC1^ΔTAK1^ with an ST2^-/-^ deletion were also subjected to papain delivery, and assessed for lung function (E; n=5-7) and histological scoring by H&E (F, G; n=7). HIC1^ΔTAK1^ versus HIC1^WT^: * p<0.05 Papain versus PBS: # p<0.05; ## p<0.01; ### p<0.01 Scale bar = 1mm

## Conclusion

MPs are used as cell-based therapeutics with limited knowledge of their normal immunomodulatory roles or how these are affected by transplantation. Here, we unveil a key immunomodulatory function of these cells, by highlighting the role of the TAK1-NF-κB signaling pathway, we suggest a way to “lock” MPs in a specific, type-2 inflammation promoting functional state that has the potential to modulate inflammatory diseases. Surprisingly, our analysis reveals a role of mesenchymal progenitors in modulating the inflammatory milieu at steady state, validating them as critical components of the immune system.

## Methods

### Animals

Housing, husbandry, and experimental protocols were conducted following institutional and national guidelines at the University of British Columbia, Canada. Mice were maintained in an enclosed and pathogen-free facility, housed in standard cages under 12-hour light–dark cycles and fed ad libitum with a standard diet. Experiments were conducted on adult female animals (8-15-week-old). Hic1-CT2:TAK1^flox/flox^ (herein Hic1^ΔTAK1^) were obtained by crossing the Hic1-CT2^7^ and the TAK1^flox/flox^ mice^23^. Hic1^ΔTAK1^ mice were then further crossed with RAG1^-/-^ mice (B6.129S7-Rag1tm1Mom/J, Jack #002216 ^44^), ST2^-/-^ (kindly gifted by Dr. G. Trinchieri, NCI), NEMO^flox/flox^ (also called IKK*γ*, kindly gifted by Dr. F. Mourkioti, UPenn).

Control mice were Hic1-CT2:TAK1^flox/WT^ (herein Hic1^WT^) or from mice with similar phenotype: RAG1^+/-^Hic1^WT^ or ST2^+/-^Hic1^WT^. For some experiments, mice also contained a Td.Tomato reporter (B6.Cg-Gt(ROSA)26Sortm9(CAG-tdTomato)Hze/J, JAX stock #007909).

Activation of the Cre recombinase was induced in controls and the various Knockouts mice by daily intraperitoneal (i.p.) injections of 3 mg/kg/day of tamoxifen (TMX, Sigma T5648) for 5 consecutive days. A washout period of 10 days was then observed to ensure no artifact effect of the TMX.

### Blood flow analysis

Mice were shaved before being placed in a restrainer. Pressure was applied to the inside of the leg and the lateral saphenous vein was punctured with a 28-gauge needle. 2-3 drops of blood were harvested in an Eppendorf tube containing 1 ml of 1X PBS 2 mM EDTA. Samples were centrifuged at 500 g for 5 minutes at 4°C and red blood cells were hemolyzed using ACK buffer (Gibco). White blood cells were then incubated with 1:1000 Fc Block (AbLab) in FACS Buffer (1X PBS 2mM EDTA 2% FBS) for 20-30 minutes before being stained for eosinophils or ILC2s (Table S4). Samples were then acquired on a LSRII (BD Aria) and data was analyzed using FlowJo™ (BD Biosciences). Eosinophils are CD45+ Lin-CD11bblow SiglecF+ ILC2 are CD45+ Lin-CD90+ CD127+ KLRG1+/-

### Blood Serum analysis

Mice were euthanized with Avertin and blood was drawn directly from the heart with a 28-gauge needle and a 1 ml syringe. Blood was left for 10 minutes at room temperature to coagulate, then spun at 12,000g for 10 minutes. Serum was collected and frozen until used for the following:

*ELISA:* IgE (BD Biosciences #555248) blood concentration was quantified by following the manufacturer instructions.

#### Mesoscale

U-PLEX Biomarker group 1 (MSD K15083K) was used to quantify EPO, GM-CSF, IFNγ, IL-1b, IL-2, IL-4, IL-5, IL-6, IL-9, IL-10, IL-12/IL-23p40, IL-12p70, IL-13, IL-15, IL-16, IL-17A/F, IL-17C, IL-17/IL-25, IL-17F, IL-21, IL-22, IL-23, IL-27p28/IL-30, IL-31, IL-33, IP-10, KC/GRO, MCP-1, MIP-1α, MIP-1β, MIP-2, MIP-3a, TNFα, VEGFA. Cytokines below the detection threshold were excluded from the analysis. Raw results are shown in Table S2.

### ELISA

Mice were euthanized with Avertin and perfused with 20ml of 1X PBS EDTA. Tissues were harvested, flash frozen into liquid nitrogen and store at -80C until use. Tissues were thawed and kept on ice. 1ml of RIPA buffer containing 1X protease inhibitor (Thermofisher #1861278) was added along with one 3mm Φ bead of tungsten (Retsch # 22.455.006). Tissues were homogenized using the TissueLyser II (Qiagen) and spun for 10min at 20,000g at +4°C. Protein concentration was determined on the supernatant by Bicinchoninic acid (BCA) (Thermofisher Pierce™)

CCL11 (Biolegend #443907) and IL-33 (Invitrogen #BMS6025) concentration were quantified by following the manufacturer instructions.

### Tissue flow analysis

Mice were euthanized with Avertin, tracheotomized with an 18-gauge catheter (BD Insyte, 150-381244) and bronchoalveolar lavage fluid (BALF) was collected as previously described ^45^. Then, mouse chest cavity was opened, and mice were perfused with 20ml of 1X PBS 2 mM EDTA. Heart, skeletal muscle, white adipose tissue, mLN were harvested. Hard tissues where processed as previously described in ^46^. Eosinophil and ILC2 staining were performed with antibodies in Table S4. mLN were dissociated and filtered through a 40 μm filter. The single cell suspension was then stained for eosinophils and ILC2 with antibodies listed in Table S4. Eosinophils are CD45+ Lin-CD11bblow CD11c-/low SiglecF+ ILC2 are CD45+ Lin-CD90+ CD127+ KLRG1+

### HIC1+ cells sorting and RNA extraction

Skeletal muscle resident HIC1+ MPs were extracted and sorted as previously described in ^47^. After sort, cells were rinsed once with 1X DEPC PBS and before being resuspended in 400ul of Trizol. RNA was precipitated overnight with isopropanol. RNA pellet was rinsed 2 time with freshly made 70% Ethanol. RNA was finally resuspended in RNAse DNAse free water containing RNAse inhibitor. Sample quality was assessed using the Agilent 2100 Bioanalyzer. RNA samples with an RNA Integrity Number > 8 were used to prepare libraries following the standard protocol for the TruSeq Stranded mRNA library kit (Illumina) on the Illumina Neoprep automated microfluidic library prep instrument. Paired-end sequencing was performed on the Illumina NextSeq 500 using the High Output Kit (75 cycle; Illumina).

### RNA-sequencing bioinformatics analysis

Illumina base call files were de-multiplexed by bcl2fastq2 (v. 2.20) on Basespace. Adaptor sequences were trimmed and low-quality reads (< 35 base pairs) were discarded. Additionally, Bowtie was used to remove read pairs that aligned against abundant sequences. Demultiplexed read sequences were then aligned to the mm10 genome reference using STAR aligner. The number of aligned reads to each annotated gene was tallied with RnaReadCounter to generate read-count matrices for all samples, which were used as inputs for downstream analyses. Bowtie, STAR, and RnaReadCounter are tools built under RNA-Seq Alignment (v 1.1.1). Downstream analyses of read-count data were performed in R (v 3.6.2). Genes with less than 2 counts per million (CPMs) in at least three samples were filtered out. Batch correction was applied using Combat-seq under the sva package (v 3.35.2; ^48^) to samples derived from different experimental dates. Filtered and batch-adjusted counts were processed and analyzed using DESeq2 (v 1.26.0; ^49^). This included count normalization, principal component analysis (PCA), and differential expression analysis. Volcano plot was generated using the EnhancedVolcano package (v 1.13.2; ^50^). Over representation analysis was performed using DAVID (v 2021 ^51,52^) with all significant genes considered to inform enriched biological processes under Gene Ontology (GO DIRECT). Bars associated with top enriched GO terms reflect log-transformed adjusted P-values. All P-values were adjusted by Benjamini-Hochberg correction.

### Data availably

RNA-seq datasets generated in the current study are available in the NCBI Gene Expression Omnibus under the accession number GSE216905

### Bone marrow analysis

Bone marrow (BM) from control and HIC1^ΔTAK1^ mice was flushed out using FACS Buffer to obtain a single cell suspension. Red blood cells were lysed with ACK buffer (Gibco) and BM suspension was stained for hematopoietic stem cells (HSC) and long-term (LT)-HSC, common myeloid progenitors (CMP), megakaryocyte/erythroid progenitors (MEP), granulocyte/monocyte progenitors (GMP), and eosinophil progenitors (EOPs) ^53,54^ with the antibodies listed in Table S4. Samples were acquired on a LSRII (BD Aria) and data was analyzed using FlowJo™ (BD Biosciences).

HSC: Lin-SCA1+ cKIT+

LT-HSC: Lin-SCA1+ cKIT+ CD48-CD150hi

CMP: Lin-SCA1-cKIT+ CD33+ CD16/32-

MEP: Lin-SCA1-cKIT+ CD34-CD16/32-

GMP: Lin-SCA1-cKIT+ CD34+ CD16/32+

EOP: Lin-SCA1-CD34+ cKITint CD125+

### T cell profiling

#### T cell activation

Spleen from control and HIC1^ΔTAK1^ mice were dissociated and red blood cells were lysed using ACK buffer (Gibco). The single cell suspension was filtered with a 40 μm filter, splenocytes were counted and incubated for 3 hours at 37°C at 1million/ml in Iscove’s modified Dulbecco’s medium (IMDM) with 20% FBS, 100 IU/ml penicillin and 10 mg/ml streptomycin, 2 mM glutamine, 25 mM HEPES, 1X nonessential amino acids, 1 mM sodium pyruvate and 0.006‰ β-mercaptoethanol containing Cell Stimulation Cocktail, PMA/Ionomycin (3ul/ml, Biolegend #423301) and brefeldin A (1X, Biolegend #420601) ^42^. Non-adherent cells were recovered and stained with surface antibody before being fixed/permeabilized (BD #554714) and stained for intracellular cytokines (Table S4).

#### Regulatory T cells

Mesenteric lymph nodes from control and HIC1^ΔTAK1^ mice were dissociated and filtered through a 40 μm filter. The single cell suspension was incubated with 1:1000 Fc Block (AbLab) in 1X FACS Buffer and stained for surface markers before being fixed/permeabilized (Thermofisher 00-5523-00) for FoxP3 staining (invitrogen 11-5773-82 Table 2).

Tregs are CD45+ CD8-CD4+ Foxp3+ CD25+/-Samples were acquired on a LSRII (BD Aria) and data was analyzed using FlowJo™ (BD Biosciences).

### CYTOF

Spleen from control and HIC1^ΔTAK1^ mice were harvested as previously described above. Splenocytes were counted and stained as follows: centrifugation steps were performed at 500 g and 4°C prior to cell fixation. All reagents were diluted according to manufacturer instructions unless stated otherwise. Cells were stained with Cell-ID™ Intercalator-Rh (Fluidigm) for 15 minutes at 37°C and washed with MaxPar® MCSB. Prior to surface staining, cells were incubated with Fc block (AbLab) for 15 minutes at 4°C and stained with a surface antibody cocktail for 30 minutes at RT (see table S4). Prior to fixation and nuclear staining, cells were washed with MaxPar® MCSB and incubated in MaxPar® Fix and Perm Buffer (Fluidigm) and Cell-ID™ Intercalator-IR (Fluidigm) overnight. Centrifugation steps were performed at 900g and 4°C post cell fixation. Cells were washed with MilliQ water and resuspended in EQ™ Four Element Calibration Beads (Fluidigm). An average of 1.5 million events were acquired for each sample at a flow rate of 45μl/minute with a CyTOF®2 mass cytometer.

Fcs files were normalized (https://github.com/nolanlab/bead-normalization) and debris, dead cells and beads were manually removed with the FlowJo gating software (BD Biosciences). Cytofkit (https://github.com/JinmiaoChenLab/cytofkit2) was used for dimensionality reduction (Uniform Manifold Approximation and Projection (UMAP)) and clustering (RPhenograph) with cytofAsinh as the transformation method. Samples were equally downsampled prior to clustering. Csv files containing mean marker expression and cluster percentages were used for further analysis and cell subset characterization. Additional pre-gating steps prior to re-clustering were manually performed to narrow in on populations of interest (e.g. CD45+Terr19-gate for removal of red blood cells). Protocol was adjusted from ^55^

### Chronic lung study

#### Chronic papain damage

Lightly anesthetized mice were administered intranasally with 10 μg of papain (Sigma, P3125) prepared in a volume of 40 μl 1X PBS at d0, d1, d14, d20, before being harvested at d21 ^42^.

#### Lung capacity

Lung function was assessed using the FlexiVent® (SCIREQ; Montréal, Canada) as previously described ^56^. Briefly, mice were anaesthetised with 100 mg/kg ketamine and 10 mg/kg xylazine, tracheotomized with an 18-gauge blunt needle and mechanically ventilated at a respiratory rate of 150 breaths/min and a tidal volume of 10 mL/kg, with a pressure limit of 30 cmH2O. Muscle paralysis was achieved using pancuronium (2 mg/kg, Sandoz; Boucherville, Canada) to prevent respiratory efforts during the measurement. The following sequence of measures was repeated three times and averaged for analysis: Deep inflation, Snapshot-150, Quick Prime-3 and Pressure/Volume-loop to obtain measures for lung capacities.

#### Histology

In order to keep the tissue integrity, lungs were inflated through a tracheal catheter at 20 cmH2O pressure with 10% formalin. The left lobe lung was ligatured, removed from the chest cavity and placed in 10% formalin for a week. Lungs were then transferred to 70% ethanol before being paraffin embedded. 4 μm sections were taken and lungs were stained with Hematoxylin and Eosin (HE) and Picrosirius (PSR) (Wax-it).

#### Scoring

Images were acquired at 10X magnification using a Nikon Eclipse Ni equipped with a device camera (Nikon Digital Sight DS-U3 for brightfield) and operated via NIS software. Lung scoring was assessed using the method described in ^43^.

#### Cytof

BAL was harvested as described above. Cells were stained with a panel of antibody (Table S5) and CYTOF was run as previously described above.

### Experimental autoimmune encephalomyelitis (EAE)

#### Procedure

EAE was performed as described in ^57^. Briefly, HIC1^ΔTAK1^ mice and their controls were anesthetized with isoflurane and immunized subcutaneously near the dorsal tail base with 200μg of rodent myelin oligodendrocyte glycoprotein (MOG35-55) over three injection sites. MOG peptide was synthesized by the Protein and Nucleic Acid Facility, Beckman Center, Stanford University. MOG was emulsified in complete Freund’s adjuvant (CFA BD B263910, Cat. DF0639-60-6) containing inactivated *Mycobacterium tuberculosis* H37Ra (6 mg/mL, BD B231141, Cat. DF3114-33-8). Mice were also co-injected with pertussis toxin (200 ng/mouse in 1X PBS, List Biological Laboratories #179A) in I.P., followed by an additional I.P. injection 48 hours after immunization.

#### Scoring

To assess disease progression, mice were weighed and scored daily until experimental endpoint or 25 days post induction, following a similar scoring as ^58^. The following system was used to assess neurological deficits: Clinical Score (CS) 0.5 – partially limp tail; CS1 – complete limp tail; CS2 – limp tail and hindlimb weakness; CS3 – hindlimb paresis; CS3.5 – single hindlimb paralysis; CS4 – complete hindlimb paralysis; CS5 – moribund.

#### Flow analysis

Mice were euthanized with i.p. injection of Avertin followed by perfusion with 30 ml of 1X PBS 2mM EDTA. Spinal cord (SP) and brain were carefully dissected away and placed in cold 1X PBS. SP and brain were digested for 20 min at 37°C in a digestion buffer containing 0.15 U/ml Collagenase D, 0.15 U/ml Dispase II, 14 ug/ml DNASe, 20 mM Cacl2 and 2% FBS). Digestion was stopped with cold 1X PBS and suspension was centrifuged at 800g for 5 minutes at 4°C. Pellet was resuspended and passed through a 40 μm filter before being centrifuged again. Pellet was then resuspended in 35% percoll (GE Healthcare, #17-0891-01) and centrifuged for 20 minutes at 800 g at room temperature. The supernatant and the intermediate layer (which contains the myelin) was carefully removed. The pellet was rinsed once with FACS Buffer and incubated with 1:1000 Fc Block (AbLab) in 1X FACS for 20 minutes before being stained for eosinophils, ILC2 and microglia (Table S4). Samples were then acquired on a LSRII (BD Aria) and data was analyzed using FlowJo™ (BD Biosciences). Microglia are defined as CD45low Ly6G/CD3/NK1/SiglecF-CD11b/CD11clow Eosinophils are defined as CD45+ Ly6G/CD3/NK1-SiglecF+ ILC2 are defined as CD45+ Lin-CD90+ CD127+

### Statistical analysis

Graph and statistical tests were performed using Prism 8 (GraphPad Software, La Jolla California, USA). A probability of <5% (p < 0.05) was considered statistically significant. Gaussian distribution was assumed. Unpaired T test, two-way Anova with Šidàk correction for multiple comparisons, or mixed effect analysis with Šidàk correction for multiple comparisons were performed. Sample size is indicated in the figure legends. Graphs are represented as mean ± standard error of the mean with individual values. Figures were assembled using Adobe Illustrator CS6 (Adobe).

## Acknowledgments

We would like to thank Dr. Foteini Mourkioti (Penn Institute for Regenerative Medicine, School of Medicine, University of Pennsylvania, Philadelphia, PA.) for the NEMO-Flox mice, and Dr. Giorgio Trinchieri (NCI) for the ST2-KO mice. We also would like to thank Andrew Johnson and Justin Wong (UBC Flow core), Takahide Murakami (genotyping), Krista Ranta, Jaspreet Rai, and Wei Yuan (Vivarium), Michael Wiliams (AbLab) as well as Lisa Kammer and Wendy Song.

## Author contribution

Theret Marine: Project design and management, experiments, data analysis and interpretation, manuscript writing, figure editing

Messing Melina: experiments, data interpretation, manuscript editing

White Zoe: experiments, data interpretation, manuscript editing

Tung Lin Wei: experiments

Rempel Lucas: experiments

Hamer Mark: experiments

Hashimoto Joshua: experiments

Li Fangfang: experiments

Brassard Julyanne: experiments

Li Yicong: experiments

Sauge Elodie: experiments

Shin Samuel: experiments

Katie Day: experiments

Manjosh Uppal: experiments

Low Marcela: preliminary data acquisition

Eisner Christine: experiments

Sato Shintaro: provided TAK1flox mice

Akira Shizuo: provided TAK1flox mice

Hughes Michael: experiments

Bernatchez Pascal: data interpretation

McNagny Kelly: data interpretation

Rossi Fabio M.V.: experimental design, data interpretation, manuscript editing and funding

## Figure legend

**Figure S1:**
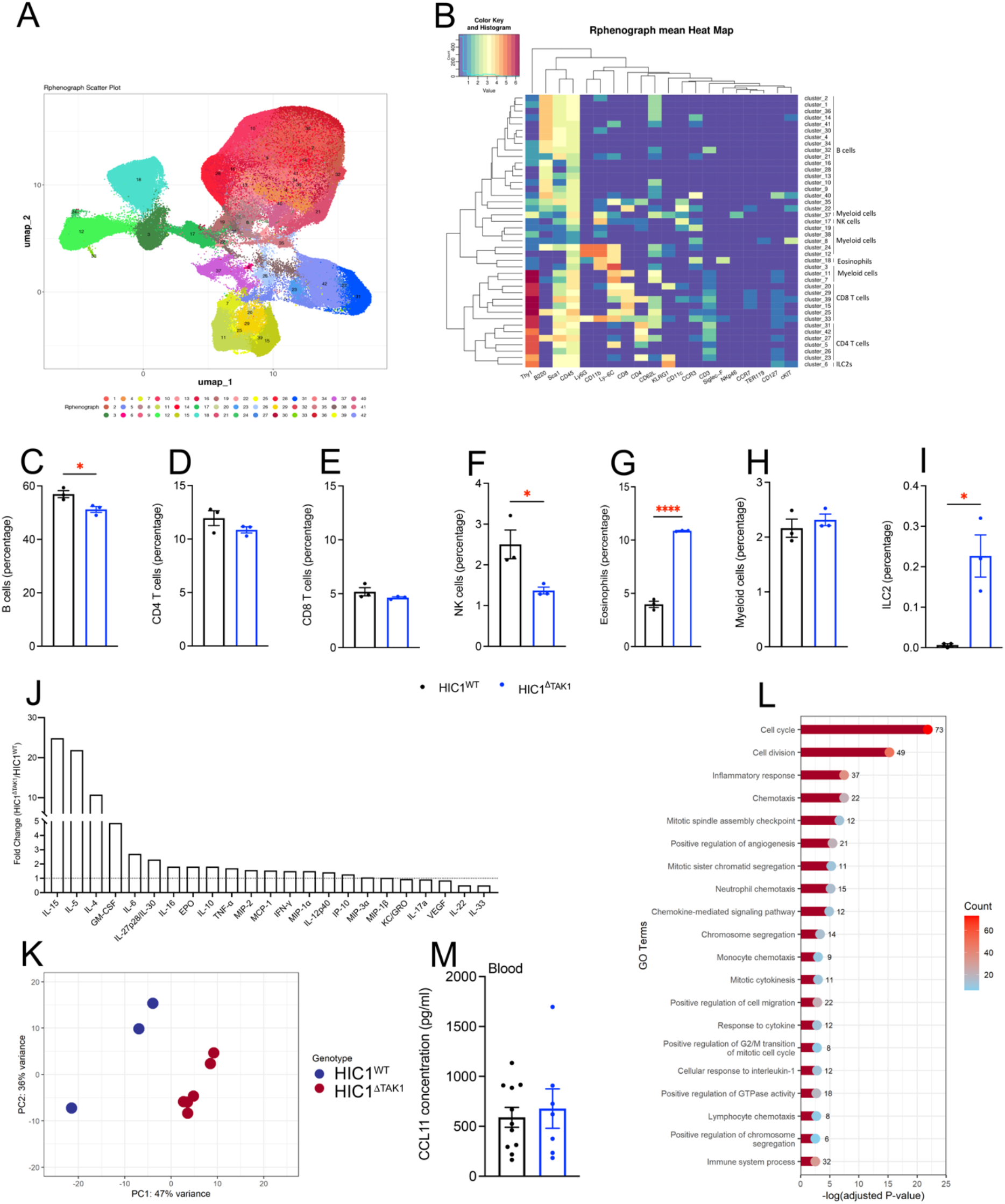
CyTOF and RNA sequencing analysis. After tamoxifen induction, splenocytes were isolated and run against a 21 antibody CyTOF panel. 42 phenotypic clusters were identified and their distributions are shown via UMAP (A) and Heat Map (B). From the 6 main populations, the percentage of B cells (C), CD4 T cells (D), CD8 T cells (E), NK cells (F), eosinophils (G), myeloid cells (H) and type 2 innate lymphoid cells (I) was calculated (n=3). Cytokine expression from HIC1^WT^ and HIC1^ΔTAK1^ serum was analysed by Mesoscale and is represented as an overall fold change (J; n=4-6). Skeletal muscle tissue resident mesenchymal progenitor from HIC1^WT^ and HIC1^ΔTAK1^ mice were sorted and subjected to RNAseq and principal component analysis (PCA) of global gene expression (K) and biological processes from GO Term analysis (L) are shown (n=3-5). CCL11 concentration from HIC1^WT^ and HIC1^ΔTAK1^ mice serum was quantified by ELISA (M; n=6-11). HIC1^ΔTAK1^ versus HIC1^WT^: * p<0.05, **** p<0.0001

**Figure S2:**
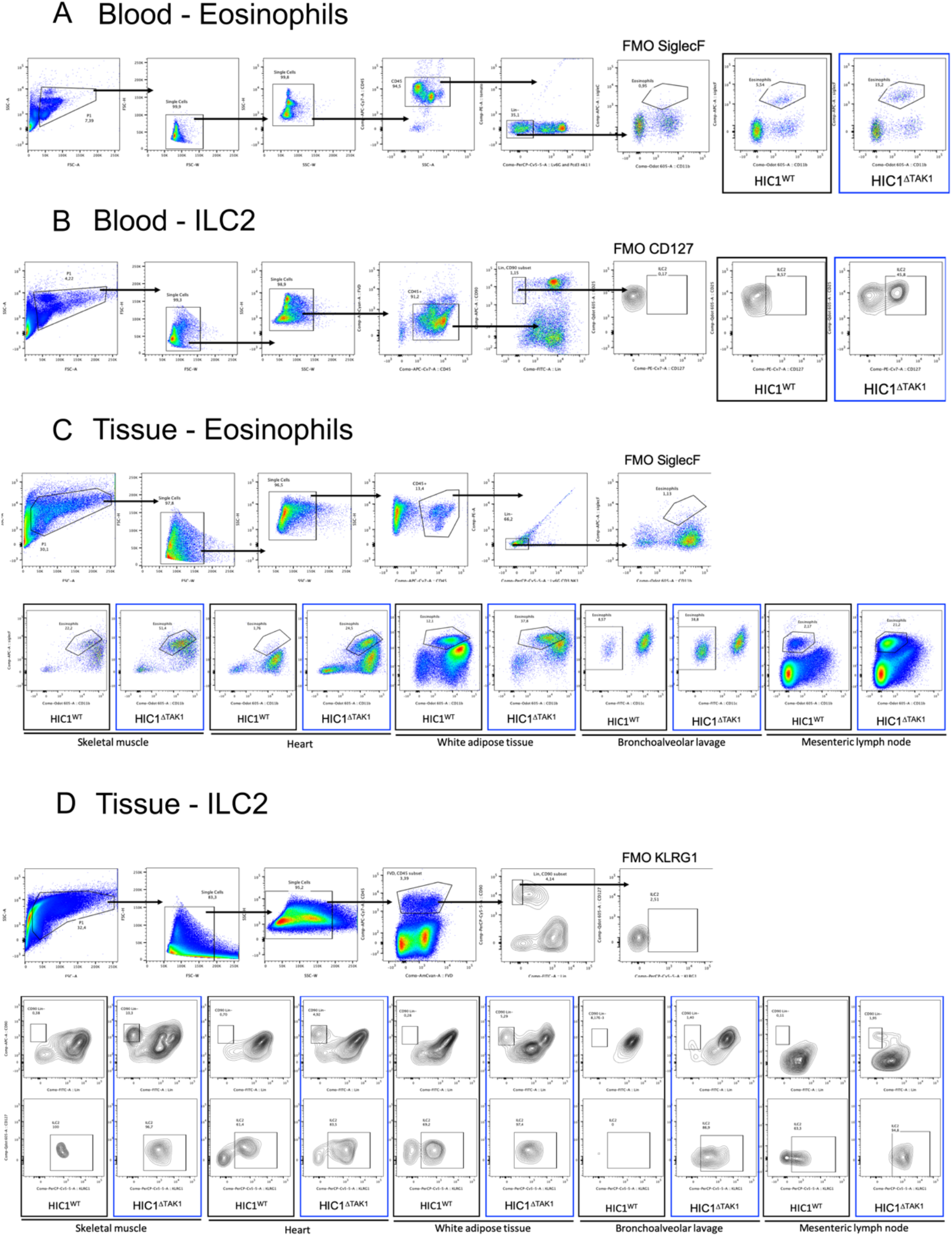
Eosinophil and type 2 innate lymphoid cell gating strategies. After tamoxifen induction, blood was freshly harvested and stained for eosinophils (CD45, NK1.1, Ly6G, CDe3, CD11b, siglecF) (A) and type 2 innate lymphoid cells (ILC2) (CD45, Lin-FITC, CD90, CD127) (B). Skm, Heart, WAT, BAL and mLN were harvested, and stained for eosinophils CD45, NK1.1, Ly6G, CDe3, CD11b, CD11c, SiglecF) (D) and ILC2 (CD45, Lin-FITC, CD90, CD127, KLRG1) (E).

**Figure S3:**
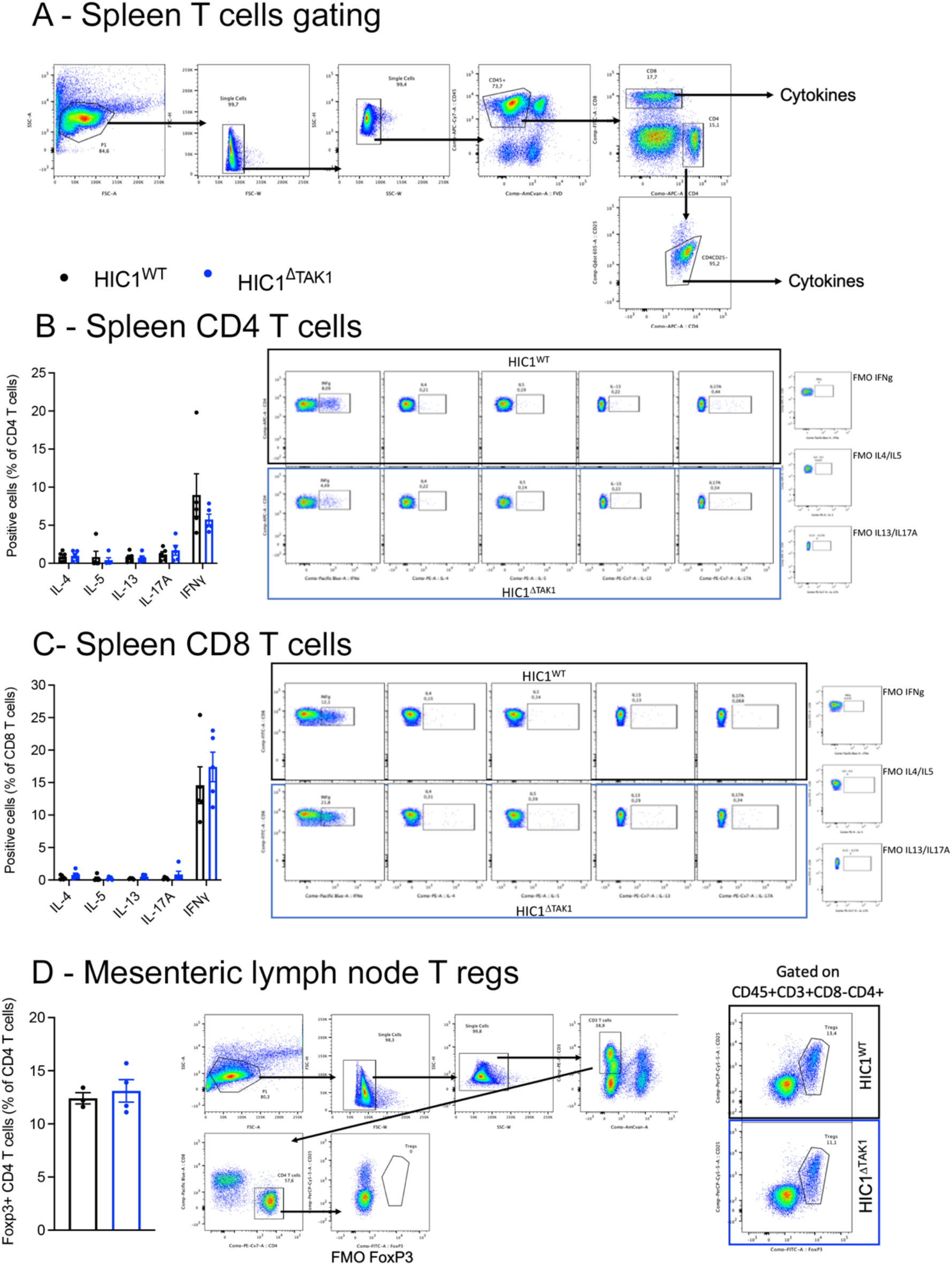
T cell profiling in HIC1^ΔTAK1^ mice. After tamoxifen induction, solenocytes were isolated and stained for CD8 and CD4 T cells (A). CD4 and CD8 T cells were subsequently stained for IFNγ, IL-4, IL-13, IL17A and IL-5 (B-C). Mesenteric lymph nodes were harvested and stained for Tregs (D).

**Figure S4:**
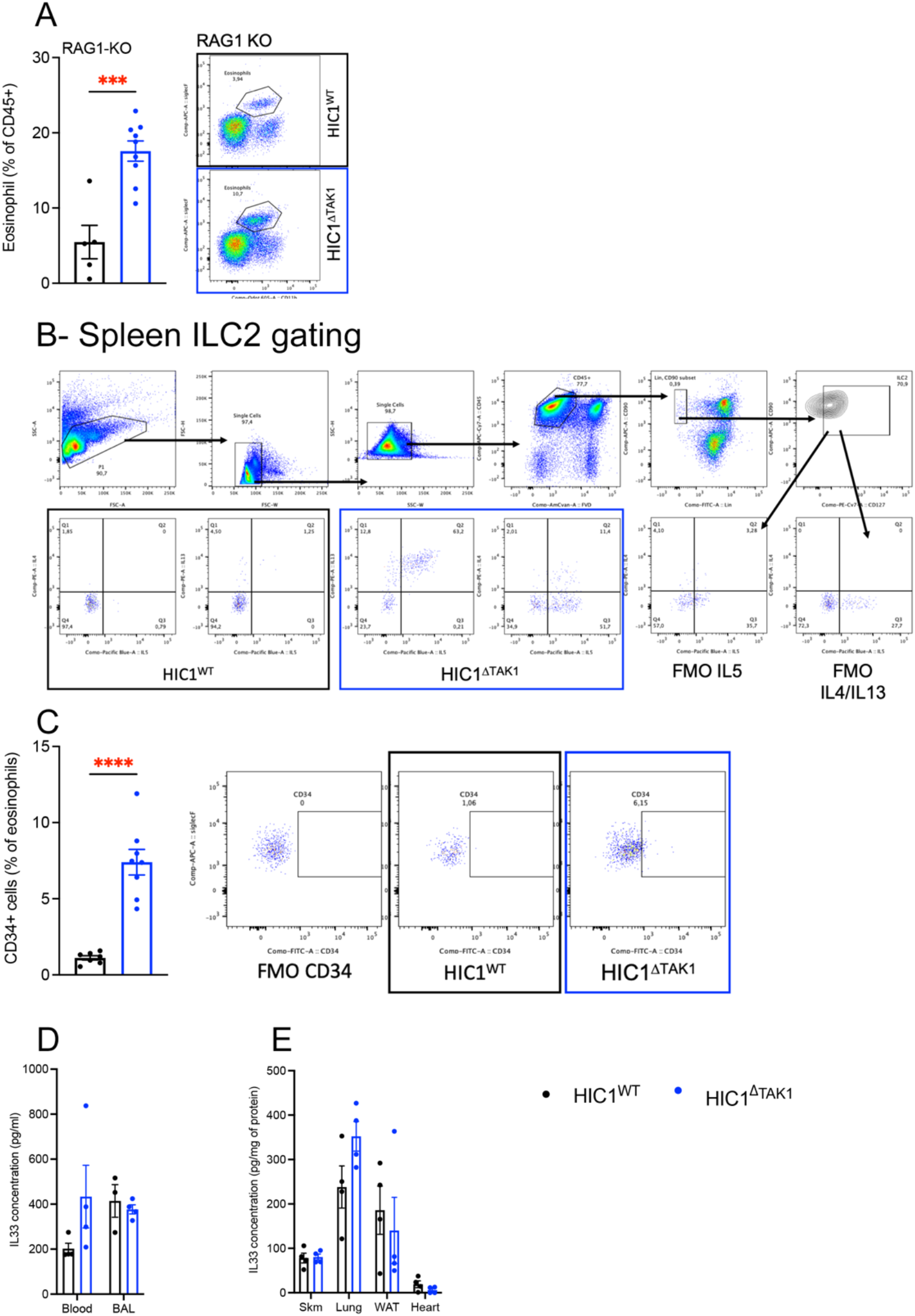
ILC2 and eosinophil profiling in HIC1^ΔTAK1^ mice. After tamoxifen induction, freshly harvested blood from RAG1-KO: HIC1^ΔWT^ and RAG1-KO: HIC1^ΔTAK1^ was stained for eosinophils (A). Splenocytes were harvested, stimulated for 3 hours and stained for type 2 innate lymphoid cells (ILC2), IL-4, IL-13, and IL-5 (B, gating strategy). Blood eosinophils from HIC1^WT^ and HIC1^ΔTAK1^ mice were stained with CD34 (C; n=7-8). IL-33 concentration was quantified in blood, BAL (D), skm, Lung, WAT, and Heart (E; n=3-4). HIC1^ΔTAK1^ versus HIC1^WT^: *** p<0.001, *** p<0.0001

**Figure S5:**
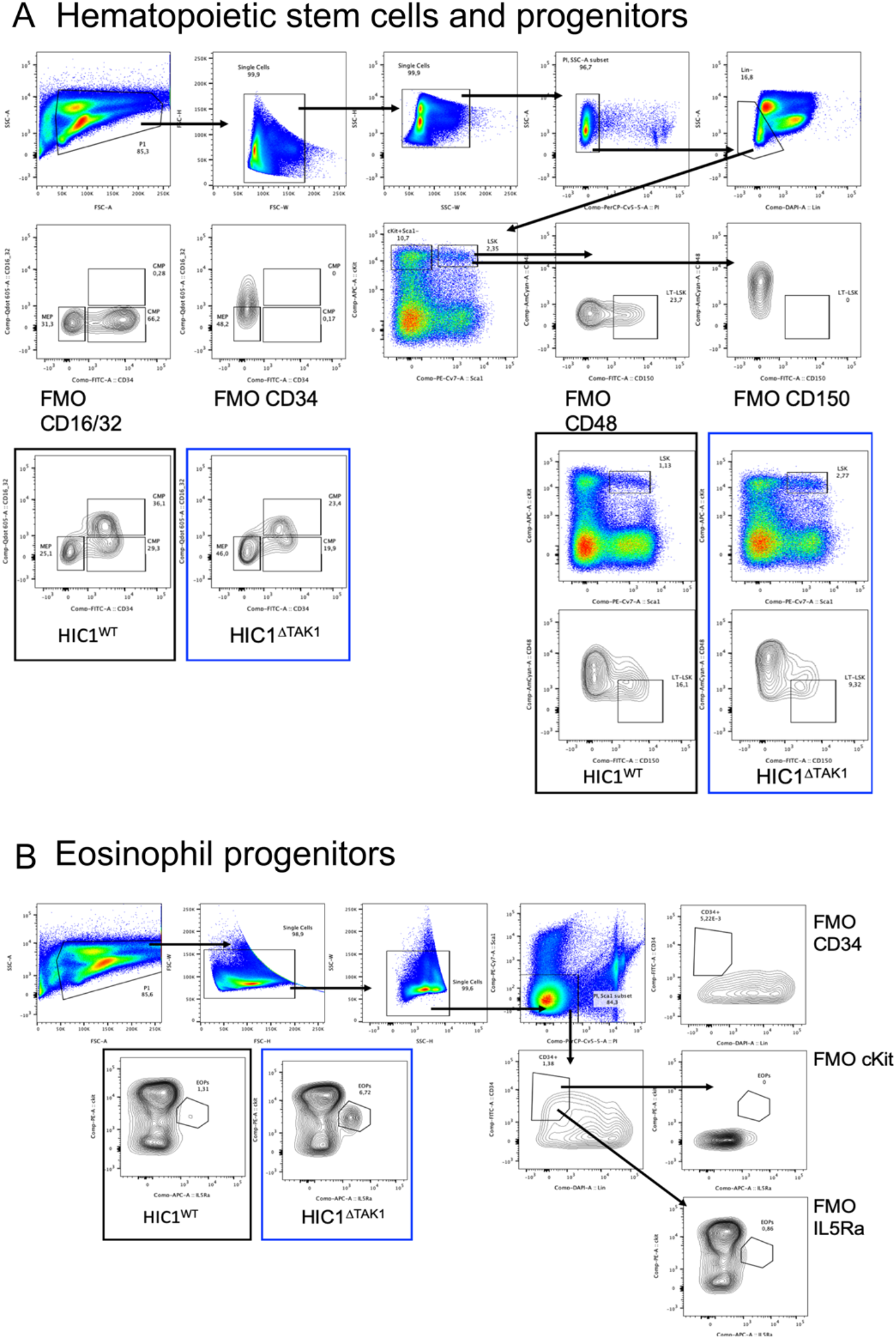
Hematopoiesis in HIC1^ΔTAK1^ mice. After tamoxifen induction, bone marrow was harvested and stained for Lin-Sca1+cKit+ (LSK) cells, Long Term LSK (LT-LSK), and various progenitors such as common myeloid progenitors (CMP), granulocyte/monocyte progenitors (GMP), megakaryocyte/erythroid progenitors (MEP) (A) and eosinophil progenitors (EOPs) (B).

**Figure S6:**
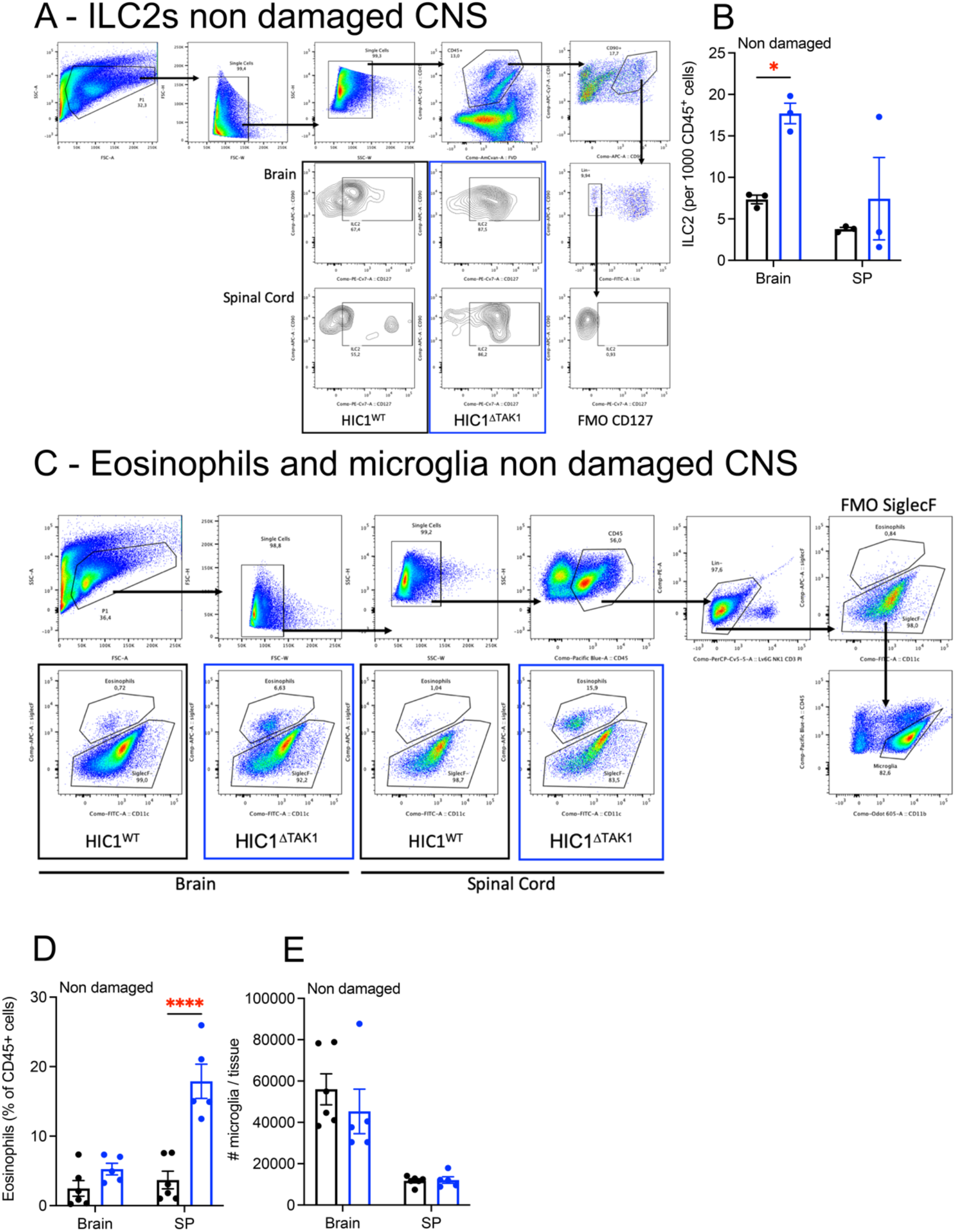
ILC2 and eosinophils in HIC1^ΔTAK1^ CNS. After tamoxifen induction, brain and spinal cord (SP) of HIC1^WT^ and HIC1^ΔTAK1^ mice were harvested and stained for ILC2 (A-B; n=3), eosinophils and microglia (C-E; n=5-6). HIC1^ΔTAK1^ versus HIC1^WT^: *** p<0.0001

**Figure S7:**
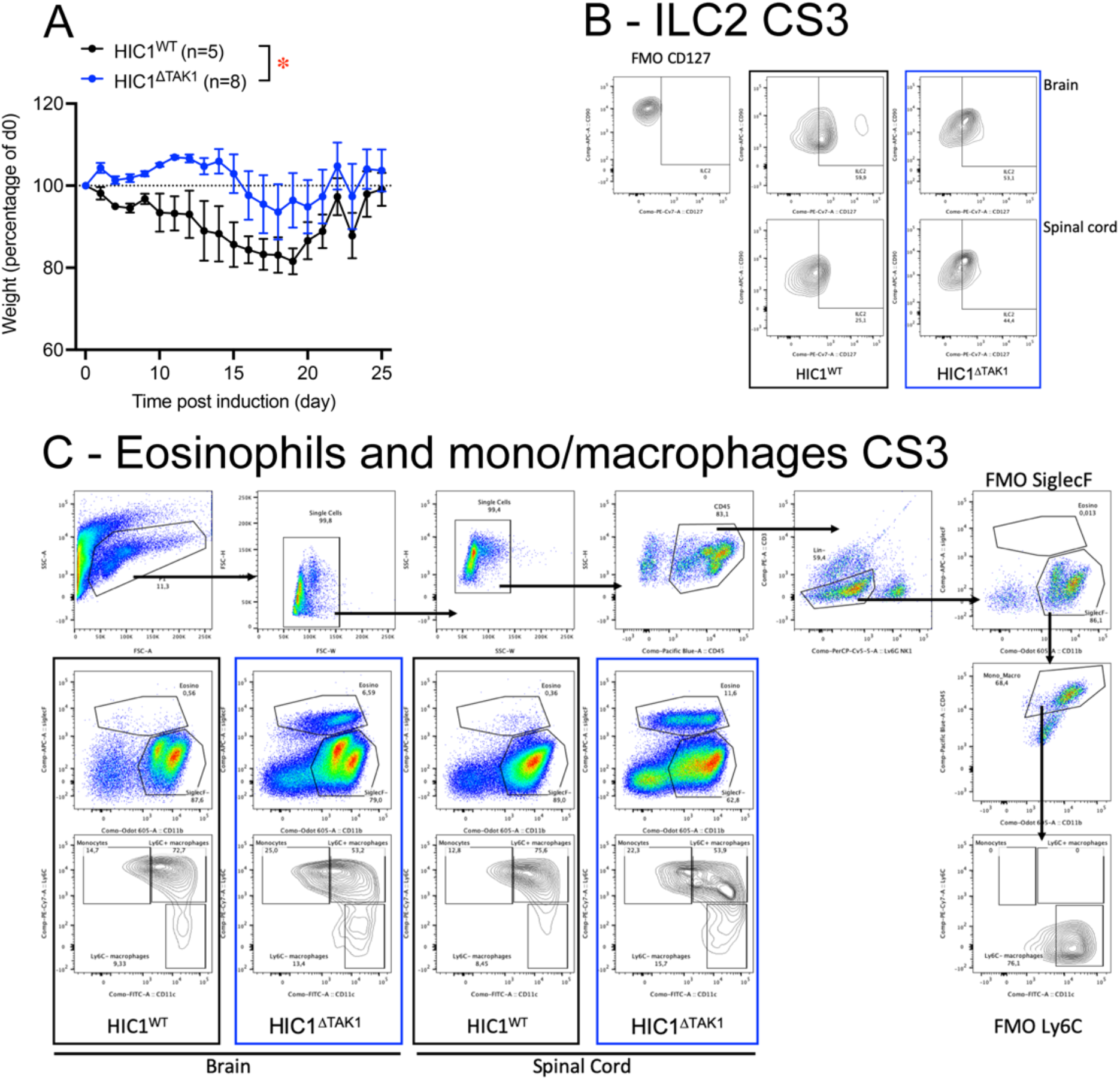
Experimental Autoimmune Encephalomyelitis. After EAE inductions, body weights of HIC1^WT^ and HIC1^ΔTAK1^ mice were collected up to 25 days post induction (A; n=5-8). The CNS was harvested when mice reached a clinical score of 3 (CS3) and stained for ILC2 (B), eosinophils and monocytes/macrophages (C). HIC1^ΔTAK1^ versus HIC1^WT^: * p<0.05

**Figure S8:**
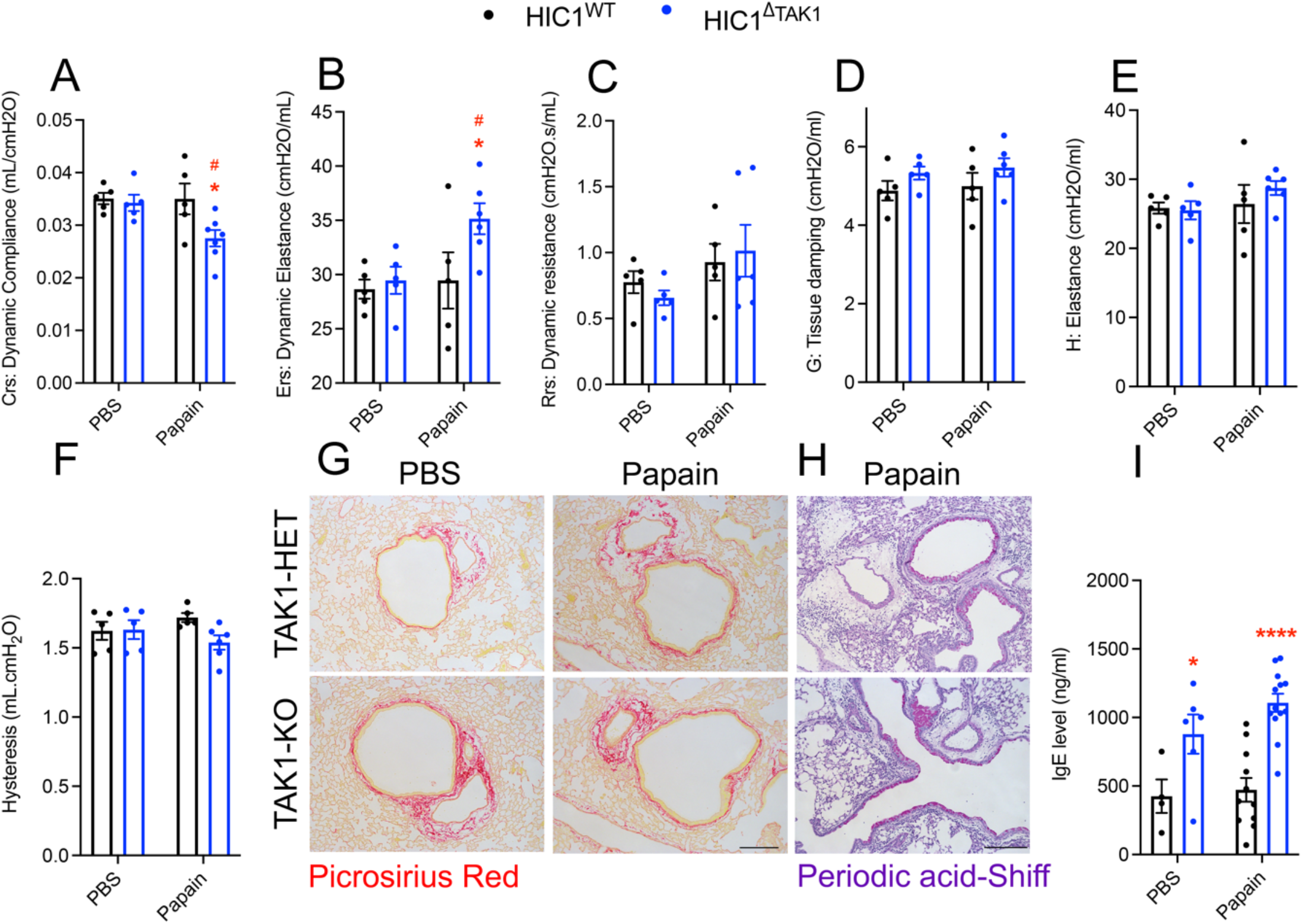
Lung function of HIC1^ΔTAK1^ mice after papain damage. After tamoxifen injections, HIC1^WT^ and HIC1^ΔTAK1^ mice were subjected to intra-nasal papain injections to induce an allergic reaction. Lung function was measured at 21 days post-injection using Flexivent® for the following parameters: compliance (A), dynamic elastance (B), resistance (C), tissue damping (D), elastance (E) and hysteresis (F) n=5-6. Lungs were harvested and stained for collagen (picrosirius red) (G) and mucus (Periodic acid Schiff) (H). Finally, serum level of IgE was quantified (I; n=4-13). HIC1^ΔTAK1^ versus HIC1^WT^: * p<0.05, ** p<0.01, *** p<0.001, **** p<0.0001 Papain versus PBS: # p<0.05 Scale bar = 200μm

**Figure S9:**
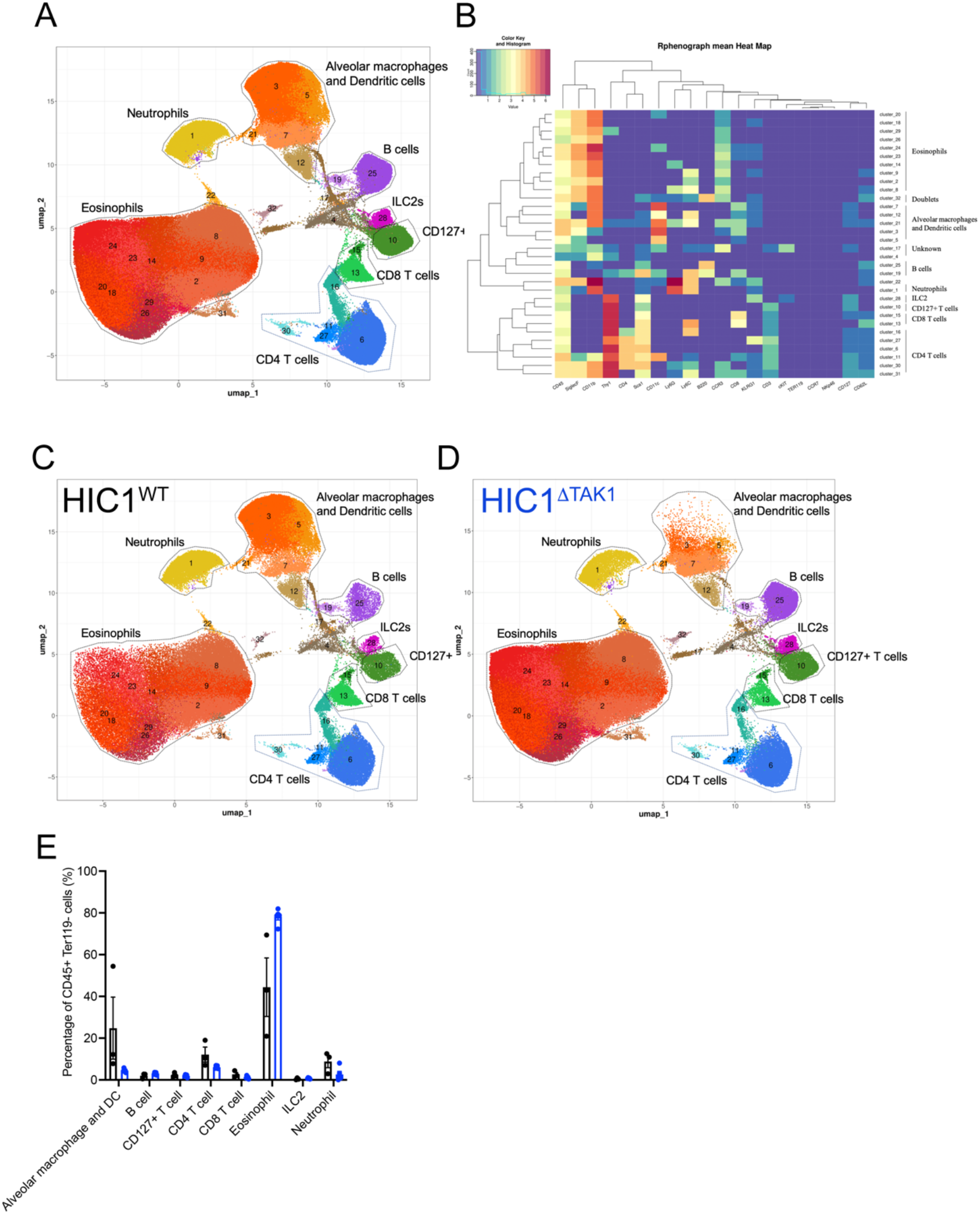
Immune cell profiling of HIC1^ΔTAK1^ mice after papain damage. After tamoxifen injections, mice were subjected to intra-nasal papain injections to induce an allergic reaction. Lungs were digested and subjected to a 21 antibody CyTOF panel. 32 clusters were identified and regrouped under B cells, CD4 T cells, CD8 T cells, CD127+ T cells, neutrophils, alveolar macrophages and dendritic cells, ILC2, and eosinophils (A-B). Percentages for each population were calculated (C-E; n=3).

**Figure S10:**
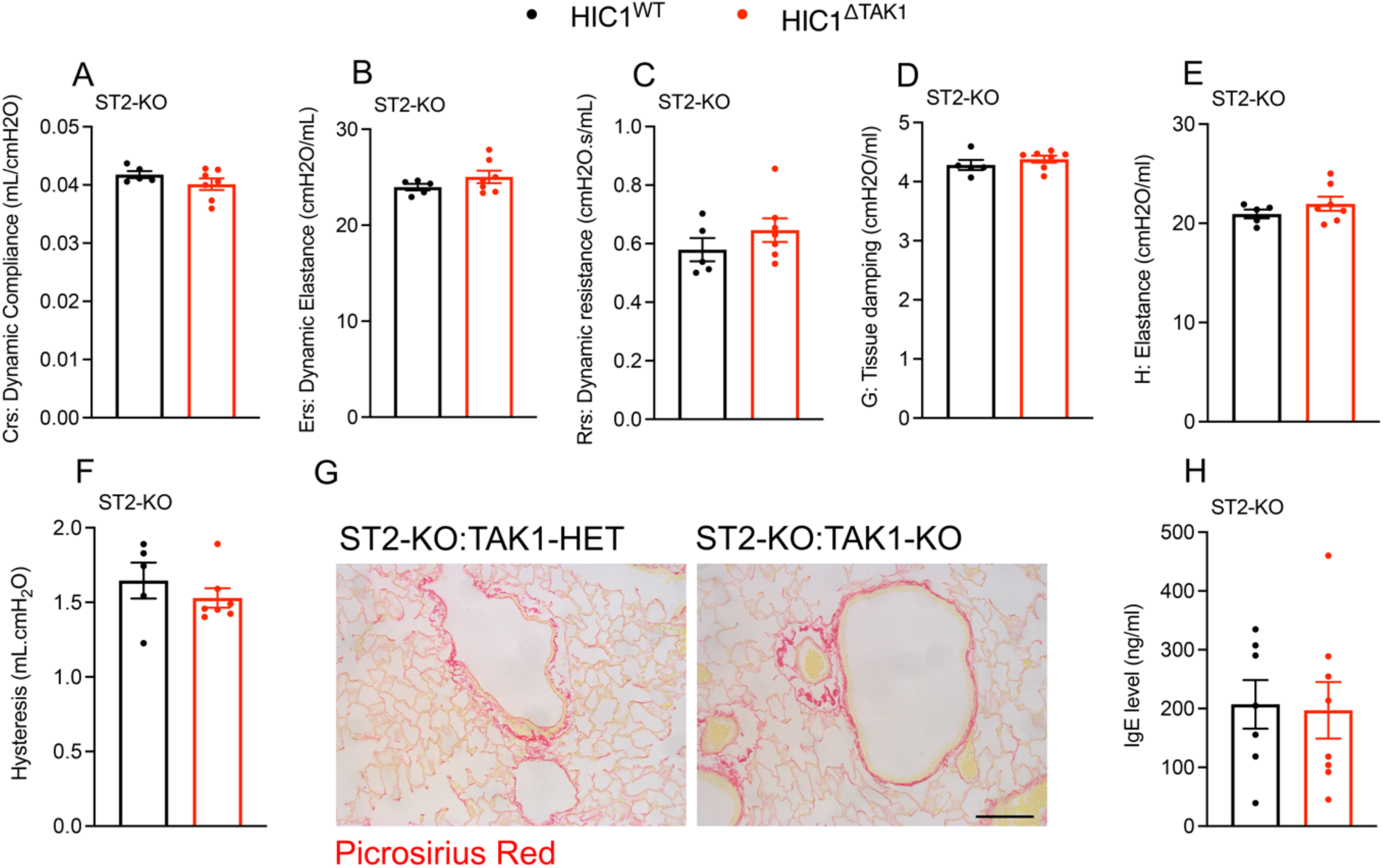
Lung function of ST2-KO:HIC1^ΔTAK1^ mice after papain damage. After tamoxifen injections, mice were subjected to intra-nasal papain injections to induce an allergic reaction. Lung function was measured 21 days post-injection using Flexivent® for the following parameters: compliance (A), dynamic elastance (B), resistance (C), tissue damping (D), elastance (E) and hysteresis (F) n=5-7. Lungs were harvested and stained for collagen (picrosirius red) (G) and serum levels of IgE were quantified (I; n=7-8). Scale bar = 200μm

## References

1. Soliman, H. et al. Multipotent stromal cells: One name, multiple identities. Cell Stem Cell 28, (2021).

2. Song, N., Scholtemeijer, M. & Shah, K. Mesenchymal Stem Cell Immunomodulation: Mechanisms and Therapeutic Potential. Trends in Pharmacological Sciences 41, (2020).

3. Lukomska, B. et al. Challenges and Controversies in Human Mesenchymal Stem Cell Therapy. Stem Cells International 2019, (2019).

4. Waterman, R. S., Tomchuck, S. L., Henkle, S. L. & Betancourt, A. M. A new mesenchymal stem cell (MSC) paradigm: Polarization into a pro-inflammatory MSC1 or an immunosuppressive MSC2 phenotype. PLoS One 5, (2010).

5. Shi, Y., Du, L., Lin, L. & Wang, Y. Tumour-associated mesenchymal stem/stromal cells: Emerging therapeutic targets. Nature Reviews Drug Discovery 16, (2016).

6. Munir, S. et al. TLR4-dependent shaping of the wound site by MSCs accelerates wound healing. EMBO Rep. 21, (2020).

7. Scott, R. W., Arostegui, M., Schweitzer, R., Rossi, F. M. V. & Underhill, T. M. Hic1 Defines Quiescent Mesenchymal Progenitor Subpopulations with Distinct Functions and Fates in Skeletal Muscle Regeneration. Cell Stem Cell 25, (2019).

8. Kastenschmidt, J. M. et al. A stromal progenitor and ILC2 niche promotes muscle eosinophilia and fibrosis-associated gene expression. Cell Rep. 35, (2021).

9. Rana, B. M. J. et al. A stromal cell niche sustains ILC2-mediated type-2 conditioning in adipose tissue. J. Exp. Med. 216, (2019).

10. Liu, O. et al. Adipose-mesenchymal stromal cells suppress experimental Sjögren syndrome by IL-33-driven expansion of ST2+ regulatory T cells. iScience 24, (2021).

11. Galleu, A. et al. Apoptosis in mesenchymal stromal cells induces in vivo recipient-mediated immunomodulation. Sci. Transl. Med. 9, (2017).

12. Prockop, D. J. Concise review: Two negative feedback loops place mesenchymal stem/stromal cells at the center of early regulators of inflammation. Stem Cells 31, (2013).

13. Romieu-Mourez, R. et al. Cytokine Modulation of TLR Expression and Activation in Mesenchymal Stromal Cells Leads to a Proinflammatory Phenotype. J. Immunol. 182, (2009).

14. Yamaguchi, K. et al. Identification of a member of the MAPKKK family as a potential mediator of TGF-β signal transduction. Science (80-.). 270, (1995).

15. Ninomiya-Tsuji, J. et al. The kinase TAK1 can activate the NIK-IκB as well as the MAP kinase cascade in the IL-1 signalling pathway. Nature 398, (1999).

16. Blonska, M. et al. TAK1 is recruited to the tumor necrosis factor-α (TNF-α) receptor 1 complex in a receptor-interacting protein (RIP)-dependent manner and cooperates with MEKK3 leading to NF-κB activation. J. Biol. Chem. 280, (2005).

17. Xu, Y. R. & Lei, C. Q. TAK1-TABs Complex: A Central Signalosome in Inflammatory Responses. Frontiers in Immunology 11, (2021).

18. Ajibade, A. A., Wang, H. Y. & Wang, R. F. Cell type-specific function of TAK1 in innate immune signaling. Trends in Immunology 34, (2013).

19. Omori, E. et al. TAK1 is a master regulator of epidermal homeostasis involving skin inflammation and apoptosis. J. Biol. Chem. 281, (2006).

20. Kajino-Sakamoto, R. et al. Enterocyte-Derived TAK1 Signaling Prevents Epithelium Apoptosis and the Development of Ileitis and Colitis. J. Immunol. 181, (2008).

21. Inagaki, M. et al. TAK1-binding protein 1, TAB1, mediates osmotic stress-induced TAK1 activation but is dispensable for TAK1-mediated cytokine signaling. J. Biol. Chem. 283, (2008).

22. Inokuchi, S. et al. Disruption of TAK1 in hepatocytes causes hepatic injury, inflammation, fibrosis, and carcinogenesis. Proc. Natl. Acad. Sci. U. S. A. 107, (2010).

23. Sato, S. et al. Essential function for the kinase TAK1 in innate and adaptive immune responses. Nat. Immunol. 6, (2005).

24. Soliman, H. et al. Pathogenic Potential of Hic1-Expressing Cardiac Stromal Progenitors. Cell Stem Cell 26, (2020).

25. Burrows, K. et al. The transcriptional repressor HIC1 regulates intestinal immune homeostasis. Mucosal Immunol. 10, (2017).

26. Brigger, D. et al. Eosinophils regulate adipose tissue inflammation and sustain physical and immunological fitness in old age. Nat. Metab. 2, (2020).

27. Stirling, R. G., Van Rensen, E. L. J., Barnes, P. J. & Chung, K. F. Interleukin-5 induces CD34+ eosinophil progenitor mobilization and eosinophil CCR3 expression in asthma. Am. J. Respir. Crit. Care Med. 164, (2001).

28. Sitkauskiene, B. et al. Regulation of Bone Marrow and Airway CD34+ Eosinophils by Interleukin-5. Am. J. Respir. Cell Mol. Biol. 30, (2004).

29. Majumdar, M. K., Thiede, M. A., Mosca, J. D., Moorman, M. & Gerson, S. L. Phenotypic and functional comparison of cultures of marrow-derived mesenchymal stem cells (MSCs) and stromal cells. J. Cell. Physiol. 176, (1998).

30. Yagi, R. et al. The transcription factor GATA3 is critical for the development of all IL-7Rα-expressing innate lymphoid cells. Immunity 40, (2014).

31. Messing, M., Jan-Abu, S. C. & McNagny, K. Group 2 innate lymphoid cells: Central players in a recurring theme of repair and regeneration. International Journal of Molecular Sciences 21, (2020).

32. Schiering, C. et al. The alarmin IL-33 promotes regulatory T-cell function in the intestine. Nature 513, (2014).

33. Stier, M. T. et al. IL-33 promotes the egress of group 2 innate lymphoid cells from the bone marrow. J. Exp. Med. 215, (2018).

34. Kato, A. Group 2 Innate Lymphoid Cells in Airway Diseases. Chest 156, (2019).

35. Kondo, Y. et al. Administration of IL-33 induces airway hyperresponsiveness and goblet cell hyperplasia in the lungs in the absence of adaptive immune system. Int. Immunol. 20, (2008).

36. Zhu, L. et al. TAK1 signaling is a potential therapeutic target for pathological angiogenesis. Angiogenesis 24, (2021).

37. Liu, T., Zhang, L., Joo, D. & Sun, S. C. NF-κB signaling in inflammation. Signal Transduction and Targeted Therapy 2, (2017).

38. Schröfelbauer, B., Polley, S., Behar, M., Ghosh, G. & Hoffmann, A. NEMO Ensures Signaling Specificity of the Pleiotropic IKKβ by Directing Its Kinase Activity toward IκBα. Mol. Cell 47, (2012).

39. Ando, D. G., Clayton, J., Kono, D., Urban, J. L. & Sercarz, E. E. Encephalitogenic T cells in the B10.PL model of experimental allergic encephalomyelitis (EAE) are of the Th-1 lymphokine subtype. Cell. Immunol. 124, (1989).

40. Langrish, C. L. et al. IL-23 drives a pathogenic T cell population that induces autoimmune inflammation. J. Exp. Med. 201, (2005).

41. Aharoni, R. et al. Specific Th2 cells accumulate in the central nervous system of mice protected against experimental autoimmune encephalomyelitis by copolymer Proc. Natl. Acad. Sci. U. S. A. 97, (2000).

42. Cait, A. et al. Microbiome-driven allergic lung inflammation is ameliorated by short-chain fatty acids. Mucosal Immunol. 11, (2018).

43. Gold, M., Marsolais, D. & Blanchet, M. R. Mouse models of allergic asthma. Methods Mol. Biol. 1220, (2015).

44. Mombaerts, P. et al. Mutations in T-cell antigen receptor genes alpha and beta block thymocyte development at different stages [published erratum appears in Nature 1992 Dec 3;360(6403):491]. Nature 360, (1992).

45. Cait, A., Messing, M., Cait, J., Canals Hernaez, D. & McNagny, K. M. Antibiotic Treatment in an Animal Model of Inflammatory Lung Disease. in Methods in Molecular Biology 2223, (2021).

46. Theret, M. et al. Elevated numbers of infiltrating eosinophils accelerate the progression of Duchenne muscular dystrophy pathology in mdx mice. Development 149, (2022).

47. Theret, M. et al. In vitro assessment of anti-fibrotic drug activity does not predict in vivo efficacy in murine models of Duchenne muscular dystrophy. Life Sci. 279, (2021).

48. Zhang, Y., Parmigiani, G. & Johnson, W. E. ComBat-seq: Batch effect adjustment for RNA-seq count data. NAR Genomics Bioinforma. 2, (2020).

49. Love, M. I., Huber, W. & Anders, S. Moderated estimation of fold change and dispersion for RNA-seq data with DESeq2. Genome Biol. 15, (2014).

50. Blighe K, Rana S & Lewis M. Publication-ready volcano plots with enhanced colouring and labeling. (2022).

51. Sherman, B. T. et al. DAVID: a web server for functional enrichment analysis and functional annotation of gene lists (2021 update). Nucleic Acids Res. 50, (2022).

52. Huang, D. W., Sherman, B. T. & Lempicki, R. A. Systematic and integrative analysis of large gene lists using DAVID bioinformatics resources. Nat. Protoc. 4, (2009).

53. Challen, G. A., Boles, N., Lin, K. K. Y. & Goodell, M. A. Mouse hematopoietic stem cell identification and analysis. Cytometry Part A 75, (2009).

54. Iwasaki, H. et al. Identification of eosinophil lineage-committed progenitors in the murine bone marrow. J. Exp. Med. 201, (2005).

55. Messing, M. et al. Prognostic peripheral blood biomarkers at ICU admission predict COVID-19 clinical outcomes. medRxiv 2022.01.31.22270208 (2022).

56. White, Z. et al. Sildenafil Prevents Marfan-Associated Emphysema and Early Pulmonary Artery Dilation in Mice. Am. J. Pathol. 189, (2019).

57. Ajami, B., Bennett, J. L., Krieger, C., Tetzlaff, W. & Rossi, F. M. V. Local self-renewal can sustain CNS microglia maintenance and function throughout adult life. Nat. Neurosci. 10, 1538–1543 (2007).

58. Dorrier, C. E. et al. CNS fibroblasts form a fibrotic scar in response to immune cell infiltration. Nat. Neurosci. 24, (2021).

