## Supplemental tables for "Mesenchymal Progenitors set the homeostatic inflammatory milieu via the TAK1-NFkB axis": Table S4_ List of antibodies used for flow cytometry.docx

- Eosinophil and monocyte staining in blood, BAL, and tissues

| **Marker** | **Colour** | **Clone** | **Reference** |
| --- | --- | --- | --- |
| CD45 | PacBlue | I3.2 | Ablab |
| Ly-6C | PE-Cy7 | HK1.4 | BioLegend |
| Ly-6G | PerCPcy5.5 | 1A8 | Biolegend |
| NK1.1 | PerCPcy5.5 | PK136 | Biolegend |
| CD3 | PerCPcy5.5 | 2C11 | Biolegend |
| CD3 | PE | 2C11 | eBioscience |
| CD170 | APC | E50-2440 | BD Bioscience |
| CD11b | BV605 | M1/70 | Biolegend |
| CD11c | FITC | N418 | Ablab |
| CD11c | FITC | N418 | invitrogen |
| CD34 | FITC | RAM34 | Invitrogen |
| PI |  | n/a |  |

- Type 2 innate lymphoid cell staining in blood and tissues

| **Marker** | **Colour** | **Clone** | **Reference** |
| --- | --- | --- | --- |
| CD45 | APC-Fire750 | 30-F11 | BioLegend |
| GR1 | FITC | RB6-8C5 | Ablab |
| NK1.1 | FITC | PK136 | Ablab |
| CD3 | FITC | KT3 | Ablab |
| CD3 | FITC | 2C11 | Ablab |
| CD4 | FITC | GK1.5 | Ablab |
| CD8a | FITC | 53.67 | Ablab |
| CD11c | FITC | N418 | Ablab |
| CD5 | FITC | 53.73 | Ablab |
| CD11b | FITC | M1/70 | Ablab |
| TCRb | FITC | H57-597 | Invitrogen |
| Ter119 | FITC | Ter119 | Ablab |
| B220 | FITC | RA-6B2 | Ablab |
| CD19 | FITC | 1D3 | Ablab |
| CD90 | APC | 53-2.1 | Invitrogen |
| CD25 | BV605 | BV605 | BioLegend |
| CD127 | PE-Cy7 | SB/199 | BD Pharmingen |
| KLRG1 | PerCPCy5.5 | 2F1 | Invitrogen |
| FVD | Efluor 506 | n/a | Invitrogen |

- Regulatory T cell staining in lymph node

| **Marker** | **Colour** | **Clone** | **Reference** |
| --- | --- | --- | --- |
| CD3 | PerCPcy5.5 | 2C11 | Biolegend |
| CD4 | PECY7 | GK1.5 | eBioscience |
| CD8 | PacBlue | 53.67 | Ablab |
| CD11b | BV605 | M1/70 | Biolegend |
| FoxP3 | FITC | FJK-16a | Invitrogen |
| CD25 | PerCPCy5.5 | PC61.5 | Invitrogen |
| FVD | Efluor 506 | n/a | Invitrogen |

- T cell staining in spleen

| **Marker** | **Colour** | **Clone** | **Reference** |
| --- | --- | --- | --- |
| CD45 | APC/Fire750 | 30-F11 | BioLegend |
| CD4 | APC | GK1.5 | Ablab |
| CD8 | FITC | 53.67 | Ablab |
| IL-5 | PacBlue | TRFK5 | Invitrogen |
| IL-4 | PE | 11B11 | Invitrogen |
| IL-13 | PE-Cy7 | 13A | Invitrogen |
| IFNg | PacBlue | XMG1.2 | eBioscience |
| IL-17A | PE | eBio17B7 | Invitrogen |
| CD25 | BV605 | PC61 | BioLegend |
| FVD | Efluor 506 | n/a | Invitrogen |

- Type 2 innate lymphoid cell staining in spleen

| **Marker** | **Colour** | **Clone** | **Reference** |
| --- | --- | --- | --- |
| CD45 | APC-Fire750 | 30-F11 | BioLegend |
| GR1 | FITC | RB6-8C5 | Ablab |
| NK1.1 | FITC | PK136 | Ablab |
| CD3 | FITC | KT3 | Ablab |
| CD3 | FITC | 2C11 | Ablab |
| CD4 | FITC | GK1.5 | Ablab |
| CD8a | FITC | 53.67 | Ablab |
| CD5 | FITC | 53.73 | Ablab |
| CD11b | FITC | M1/70 | Ablab |
| TCRb | FITC | H57-597 | Invitrogen |
| Ter119 | FITC | Ter119 | Ablab |
| B220 | FITC | RA-6B2 | Ablab |
| CD19 | FITC | 1D3 | Ablab |
| CD90 | APC | 53-2.1 | Invitrogen |
| CD127 | PE-Cy7 | SB/199 | BD Pharmingen |
| IL-5 | PacBlue | TRFK5 | Invitrogen |
| IL-4 | PE | 11B11 | Invitrogen |
| IL-13 | PE | eBio13A | Invitrogen |
| FVD | Efluor 506 | n/a | Invitrogen |

- Hematopoietic stem cell staining in bone marrow

| **Marker** | **Colour** | **Clone** | **Reference** |
| --- | --- | --- | --- |
| CD3e | PacBlue | KT3 | Ablab |
| Ter119 | PacBlue | ter119 | eBioscience |
| B220 | PacBlue | RA3-6B2 | eBioscience |
| GR1 | PacBlue | RB6-8C5 | eBioscience |
| CD11b | PacBlue | M1/70 | eBioscience |
| cKit | APC | 2B8 | eBioscience |
| CD150 | Al488 | TC15-12F12.2 | BioLegend |
| CD48 | BV510 | HM48-1 | BD Pharmingen |
| SCA-1 | PEcy7 | D7 | eBioscience |
| PI |  | n/a |  |

- Progenitor staining in bone marrow

| **Marker** | **Colour** | **Clone** | **Reference** |
| --- | --- | --- | --- |
| CD3e | PacBlue | KT3 | Ablab |
| Ter119 | PacBlue | ter119 | eBioscience |
| B220 | PacBlue | RA3-6B2 | eBioscience |
| GR1 | PacBlue | RB6-8C5 | eBioscience |
| CD11b | PacBlue | M1/70 | eBioscience |
| cKit | PECy7 | 2B8 | eBioscience |
| CD34 | FITC | RAM34 | eBioscience |
| CD16/32 | BV605 | 2.4G2 | BD Bioscience |
| SCA-1 | PERCPCy5.5 | D7 | eBioscience |
| PI |  | n/a |  |

- Eosinophil progenitor staining in bone marrow

| **Marker** | **Colour** | **Clone** | **Reference** |
| --- | --- | --- | --- |
| CD3e | PacBlue | KT3 | Ablab |
| CD4 | PacBlue | GK1.5 | eBioscience |
| CD8 | PacBlue | 53.67 | eBioscience |
| B220 | PacBlue | RA3-6B2 | eBioscience |
| GR1 | PacBlue | RB6-8C5 | eBioscience |
| CD19 | PacBlue | 1D3 | eBioscience |
| SCA-1 | PECy7 | D7 | eBioscience |
| cKit | PE | eB8 | eBioscience |
| CD34 | FITC | RAM34 | eBioscience |
| IL5Ra | APC | DIH37 | BioLegend |
| CD170 | BV510 | E50-2440 | DB Bioscience |
| PI |  |  |  |
