## Supplemental tables for "Mesenchymal Progenitors set the homeostatic inflammatory milieu via the TAK1-NFkB axis": Table S5_ Cytof Antibody panel.docx

| **Marker** | **Metal** | **Clone** |
| --- | --- | --- |
| CD45 | 89Y | 30-F11 |
| CD3 | 115In | KT3 |
| CD4 | 141 Pr | RM4-5 |
| NKp46 | 142 Nd | 29A1.4 |
| CD127 | 144 Nd | A7R34 |
| CD117 | 147 Sm | 2B8 |
| CD170 | 150 Nd | 1RNM44N |
| SCA-1 | 152 Nd | D7 |
| Ly-6C | 153 Eu | HK1.4 |
| CD62L | 154 Sm | MEL-14 |
| KLRG1 | 155 Gd | 2F1 |
| CD11b | 156 Gs | M1/70 |
| B220 | 160 Gd | RA3-6B2 |
| Ter119 | 164 Dy | TER-119 |
| Ly-6G | 165 Ho | 1A8 |
| CCR3 | 166 Er | JO73E5 |
| TCRb | 167 Er | H57-597 |
| CD90 | 169 Tm | 53-2.1 |
| CD11c | 171 Yb | N418 |
| CD8 | 172 Yb | 53-6.7 |
| CCR7 | 173 Yb | 4B12 |

Table S4: Antibody list for CYTOF
